# Spinal VGLUT3 lineage neurons drive visceral mechanical allodynia but not visceromotor reflexes

**DOI:** 10.1101/2022.09.07.507044

**Authors:** Lu Qi, Shing-Hong Lin, Qiufu Ma

## Abstract

Visceral pain is among the most prevalent and bothersome forms of chronic pain, but their transmission in the spinal cord is still poorly understood. Here we used a focal colorectal distention (fCRD) method to drive visceromotor responses (VMRs) plus affective pain-indicative aversive learning. We first found that spinal CCK neurons were necessary for noxious fCRD to drive both VMRs and aversion. We next showed that spinal VGLUT3 neurons mediate affective visceral allodynia, whose ablation caused loss of aversion evoked by low-intensity fCRD in mice with gastrointestinal (GI) inflammation or spinal circuit disinhibition. Importantly, these neurons are dispensable for driving VMRs. Anatomically, VGLUT3 neurons send projection to the parabrachial nuclei, whose photoactivation sufficiently generated aversion in mice with GI inflammation. Our studies suggest the presence of different spinal substrates that transmit nociceptive versus affective dimensions of visceral sensory information.

## INTRODUCTION

Pain from visceral organs and other deep tissues is a prevalent clinical problem but has been greatly understudied in comparison with pain evoked from the skin (Brierley et al., 2018; Cervero, 2009; Christianson and Davis, 2010; Fuentes and Christianson, 2018; Gebhart and Bielefeldt, 2016; Grundy et al., 2019; Johnson et al., 2020; West and McVey Neufeld, 2021; Yuan and Greenwood-Van Meerveld, 2021). The aim of this study was to identify spinal substrates transmitting visceral sensory information, and along the way, we wanted to revisit surrogate assays used to measure the affective component of visceral pain. With the development of genetic and viral tools, the past decade has seen enormous progress in characterizing spinal excitatory and inhibitory neurons that transmit or gate cutaneous pain, both under naïve and pathological conditions (Duan et al., 2018; Koch et al., 2018; Moehring et al., 2018; Peirs et al., 2020; Wang et al., 2022). One outcome from these studies is the revelation of distinct spinal substrates that drive exteroceptive reflexive-defensive reactions to external threats versus interoceptive self-caring responses (for example, persistent licking) to actual body injuries that drive affective tonic pain (Choi et al., 2020; Huang et al., 2019; Ma, 2022; Peirs et al., 2020; Wang et al., 2022). As a result of this circuit-level segregation, a selective loss of cutaneous affective pain may not be detected via assays that only measure first-line reflexive-defensive reactions (Ma, 2022; Mogil, 2020). Meanwhile, despite great clinical relevance and unmet medical need, modern genetic and viral tools have rarely been used to characterize spinal circuits transmitting visceral sensory information, even though electrophysiological, molecular and behavioral studies have revealed sensitization in spinal neurons following visceral inflammation or stress (Defaye et al., 2022; Farrell et al., 2017; Gamboa-Esteves et al., 2004; Harrington et al., 2012; Honore et al., 2002; Lai et al., 2011; Li et al., 2020; Long et al., 2022; Nishida et al., 2022; Traub, 2000; Wang et al., 2005; Wu et al., 2022; Zhang et al., 2013). As such, it is still poorly understood how nociceptive and affective dimensions of visceral sensory information are transmitted through the spinal cord.

In 1988, Ness and Gebhart introduced two assays to study nociception and aversion evoked by colorectal distention (CRD) (Ness and Gebhart, 1988; Ness et al., 1991), a stimulus producing pain percepts in humans (Ritchie, 1973; Whitehead et al., 1990). One measures nociception via electromyographic recording of abdominal muscle contractions, referred to as visceromotor responses (VMRs). The other measures aversion by using the real-time step-down aversive learning assay (Ness and Gebhart, 1988; Ness et al., 1991). Aversion and distress can also be measured via the two-chamber conditioned place avoidance assay or recording post-stimulation ultrasound vocalization (Ji et al., 2015; Yan et al., 2012). While aversive learning and distressful vocalization might reflect affective experience, VMRs have been widely used as the surrogate assay to measure visceral pain (Cao et al., 2021; Christianson et al., 2010; Crock et al., 2012; DeBerry et al., 2015; Hu et al., 2020; Johnson et al., 2020; Johnson et al., 2018; Kiyatkin et al., 2013; Lai et al., 2011; Laird et al., 2001; Laird et al., 2000; Laird et al., 2002; Louwies et al., 2021; Makadia et al., 2018; Ness et al., 2018; Traub et al., 2008). Nonetheless, the gastrointestinal (GI) tract does represent a semi-external world and CRD-evoked abdominal muscle contraction represents a defensive reaction that increases intraperitoneal pressure and helps get rid of harmful objects along the GI tract (Drake et al., 2015; Feng and Guo, 2020). Furthermore, VMRs were preserved in decerebrated animals (Ness and Gebhart, 1988), whereas in humans and animals, cortical structures such as the anterior cingulate cortex (ACC) are crucial for the experience of affective pain and emotional distress (Ma, 2022; Price, 2002; Xiao and Zhang, 2018). Consistently, lesions of ACC led to a loss of affective inflammatory visceral pain measured via conditioned place avoidance, without affecting sensitized VMRs (Yan et al., 2012), and conversely, some drugs that blocked VWRs failed to treat visceral pain in humans (Blackshaw, 2012). ACC, however, resides at a very downstream step in processing affective visceral sensory information. To date, it remains entirely unknown if there are distinct spinal substrates that drive affective pain versus defensive VMRs.

Here we showed that spinal VGLUT3 lineage neurons, marked by transient developmental expression of the vesicular glutamate transporter 3 gene (*Vglut*3) (Cheng et al., 2017; Peirs et al., 2015), are required to drive aversion evoked by low-intensity CRD in mice with GI tract inflammation or with spinal circuit disinhibition. Strikingly, ablation of these neurons did not affect CRD-evoked sensitized VMRs. We also identified spinal CCK neurons required to drive both VMRs and aversion evoked by noxious fCRD under naïve conditions. Our studies suggest the necessity of using multiple physiological and behavioral assays to capture distinct dimensions of visceral sensory information.

## RESULTS

### A focal CRD method for driving VMRs and aversion

Measurement of CRD-evoked visceral nociception traditionally requires colonic insertion of a long balloon and the use of a barostat to control distension (Christianson and Gebhart, 2007). The high cost of the barostat instrument might, however, have discouraged investigators from getting into the visceral pain field. Here we modified a cost-effective focal CRD (fCRD) method first developed by Annahazi et al. (Annahazi et al., 2012). The GI tract is dually innervated by spinal and vagal sensory neurons residing in dorsal root ganglia (DRGs) and nodose ganglia, respectively, and activation of vagal afferents is capable of producing aversive feeling as well (Furuta et al., 2012; Gebhart and Bielefeldt, 2016; Lamb et al., 2003; Meerschaert et al., 2020; Randich and Gebhart, 1992). Our goal was to study spinal substrates transmitting aversive visceral information. We accordingly performed retrograde labeling (Figure S1), aimed to identify the fCRD spot innervated predominantly by spinal afferents. Placing the Fluorogold tracer in the rectum (0.5-1 cm from the anus, Figure S1A), at either the backside or the frontside, led to neuronal labeling in lumbosacral (L6-S2) DRGs, with little labeling in nodose ganglia (Figures S1A). For retrograde labeling from the colon (3.5-4.0 cm from the anus), backside injection to spots adjacent to mesenteric attachment led to retrograde labeling in nodose ganglia, thoracolumbar (T10-L2) DRGs and L6-S1 DRGs, whereas frontside injection produced very few labeled cells in the S1 DRG (Figure S1B). This differential innervation along the back-front axis in the colon versus the rectum is consistent with previous electrophysiological recordings (Brierley et al., 2018; Feng and Guo, 2020). The lack of vagal innervation in the rectum is also consistent with progressive reduction of vagal afferent innervations along the proximal-distal axis of the gastrointestinal (GI) tract (Wang and Powley, 2000), but conflicts with other studies (Christianson et al., 2006 ; Meerschaert et al., 2020; Robinson et al., 2004) (see discussion in Figure S1A). We accordingly chose to perform fCRD at distal rectum to help study spinal transmission of visceral information (but see the Discussion section on potential involvement of vagal afferents under pathological conditions). Homemade latex balloons were used, with different degrees of fCRD achieved by injecting different volumes of water (0-100μl) (Figure S2A).

We next found that fCRD in naïve mice was able to drive visceromotor responses (VMRs) detected through electromyographic (EMG) recordings of abdominal muscle contractions. For both males and females, 25μl fCRD did not produce any responses, 50μl fCRD started to produce low-level EMG activity, and both 75μl fCRD and100μl fCRD produced robust EMG activity compared with sham 0μl fCRD (Figure 1A and 1B). In overall, females appeared to show stronger responses (Figure 1A and 1B), consistent with previous reports (Holdcroft et al., 2000; Ji et al., 2012). To determine if fCRD sufficiently produced affective pain, we performed the real-time step-down assay (Ness and Gebhart, 1988). Whenever mice stepped down from a flipped petri dish to the floor, fCRD was delivered (Figure 1C). We found that 75μl fCRD was sufficient to produce a negative teaching signal, and after 4-6 trials, most mice learned to stay onto the dish, for both males and females (Figure1D and 1E). In contrast, 25μl fCRD did not cause any increase in aversive learning compared with sham 0μl fCRD (Figure1D and 1E). Thus 25μl fCRD and 75μl fCRD represent innocuous and noxious stimuli, respectively. Consistent with minimal vagal afferent innervation in the distal rectum, bilateral transection of subdiaphragmatic vagal nerves did not affect 75μl fCRD-evoked aversion (Figure S2B). In other words, spinal afferents were sufficient to drive nociception and aversion in response to rectal fCRD, although caution is needed to interpret results from vagotomy (see Figure S2B).

**Figure 1.**
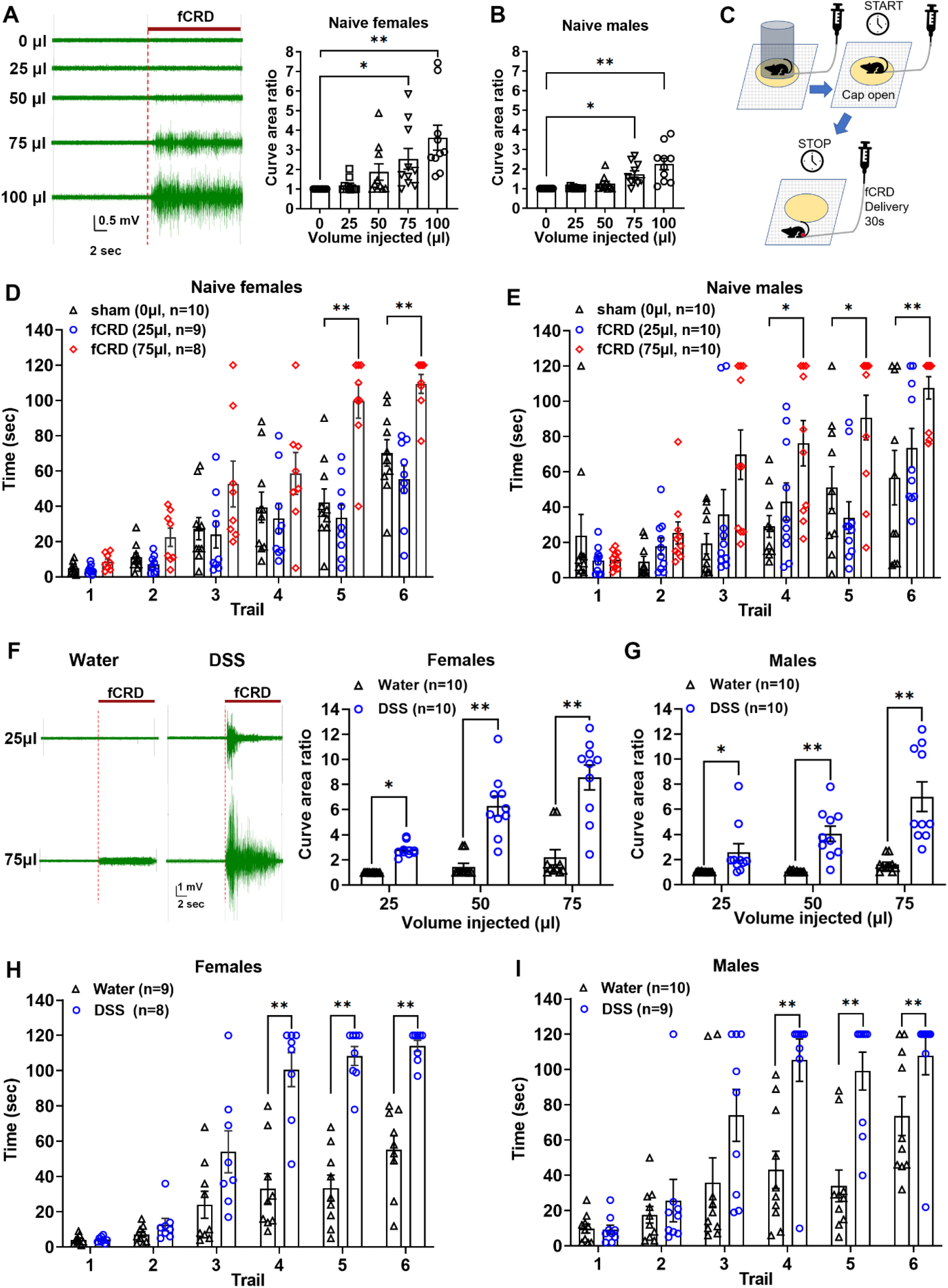
fCRD-evoked VMRs and aversion under acute and DSS-treated conditions. (A, B) fCRD-evoked VMRs in naïve females (A) and males (B). Representative traces of electromyographic (EMG) activity indicating abdominal muscle contractions evoked by 0-100μl fCRD (B, left). Volume-dependent increase of EMG activity compared with 0μl fCRD, for both females and males (one-way ANOVA, n = 10 per group; females: *F*_4,45_ = 6.52, p < 0.01; males: *F*_4,45_ = 11.00, p < 0.01; post-hoc Holm-Sidak test: *p < 0.05, **p < 0.01). (C) Schematic illustration of the step-down aversive learning assay (for details, see methods). (D, E) Using the step-down assay to assess aversive learning evoked by 25 or 75μl fCRD compared with 0μl fCRD, in both naïve females and males (two-way ANOVA; females: n = 8, 9, 10, *F*_10,120_ = 3.77, p < 0.01; males: n = 10 per group, *F*_10,135_ = 2.67, p < 0.01; post-hoc Holm-Sidak test revealed a difference of 75μl vs 0μl: *p < 0.05, **p < 0.01; no difference between 25μl vs 0μl, males: p = 0.23-0.97; females: p = 0.12-0.74). (F, G) Sensitized VMRs in mice with colitis. Representative EMG traces induced by 25 or 75 μl fCRD in females (F, left). Increased fCRD-evoked EMG activity in DSS-drinking mice compared with water-drinking mice, for both females and males (two-way ANOVA, n = 10 per group; females, *F*_2,36_ = 12.37, p < 0.01; males: *F*_2,36_ = 12.26, p < 0.01; post-hoc Holm-Sidak test: *p < 0.05, **p < 0.01). (H, I) 25μl fCRD evoked aversive learning in DSS-drinking mice compared with minimal learning in water-drinking mice (two-way ANOVA; females: n = 8, 9, *F*_5,75_ = 13.48, p < 0.01; males: n = 9, 10, *F*_5,85_ = 4.14, p < 0.01; post-hoc Holm-Sidak test: **p < 0.01). Data are shown as mean ± SEM.

We next explored if this fCRD can drive sensitized VMRs and aversion under inflammatory conditions, using a colitis model induced by drinking 3% dextran sulfate sodium (DSS) for seven days (Figure S2C) (Chassaing et al., 2014). Under this inflammatory condition, 25μl fCRD gained ability to produce VMRs, and both 50μl and 75μl fCRD produced exaggerated VMRs in comparison with naïve mice (Figure 1F and 1G). Furthermore, the step-down assay showed that 25μl fCRD now gained ability to produce negative teaching signals in DSS-treated males and females, as indicated by robust avoidance after 4-6 trials, in contrast to minimal learning in naïve mice (Figure 1H and 1I). These results indicate that 25μl rectal fCRD, which is innocuous under naïve conditions, can sufficiently drive visceromotor reflexes and aversion following colitis induction.

### Spinal CCK neurons transmitted acute noxious visceral mechanical information

While this study focused on spinal substrates transmitting inflammatory visceral pain (see below), our pilot screening also identified spinal neurons transmitting acute noxious visceral mechanical information (Figures S3-S5). For these acute studies, we investigated three groups of spinal neurons marked by developmental expression of VGLUT3-Cre, SOM-Cre or CCK-Cre, in which the *Cre* recombinase was separately driven from the *Vglut3, somatostatin*, or *cholecystokinin* gene locus (Abraira et al., 2017; Cheng et al., 2017; Duan et al., 2014; Gatto et al., 2021; Liu et al., 2018; Peirs et al., 2021; Peirs et al., 2015). We then used the intersectional genetic strategy (Figure S3A) to create *Vglut3*^*Lbx1*^-*DTR, SOM*^*Lbx1*^-*DTR*, and *CCK*^*Lbx1*^-*DTR* mice, in which the diphtheria toxin (DT) receptor DTR was expressed selectively in spinal VGLUT3^Lbx1^, SOM^Lbx1^ or CCK^Lbx1^ neurons defined by coexpression of VGLUT3-Cre, SOM-Cre or CCK-Cre plus Lbx1-Flpo, with the Lbx1-Flpo expression restricted to relay somatic sensory neurons in the spinal cord and hindbrain trigeminal nuclei, as reported previously (Bourane et al., 2015; Cheng et al., 2017; Duan et al., 2014; Gatto et al., 2021). These mice additionally carried a Cre-dependent tdTomato reporter allele, such that all *Vglut3*^*Cre*^, SOM^Cre^ or CCK^Cre^ lineage cells were marked with tdTomato (Figure S3). Repeated DT injections led to ablation of spinal VGLUT3^Lbx1^, SOM^Lbx1^ or CCK^Lbx1^ neurons (Figure S3B-D). We referred to these ablation mice as VGLUT3^Lbx1^-Abl, SOM^Lbx1^-Abl and CCK^Lbx1^-Abl, respectively. Littermates lacking DTR expression but receiving the same DT injections were used as control.

Previous cutaneous nociception studies showed that spinal VGLUT3^Lbx1^ neurons drive withdrawal responses to light punctate noxious mechanical stimuli applied to the hindpaw via von Frey filaments (Cheng et al., 2017; Peirs et al., 2015). Spinal SOM^Lbx1^ neurons play a broader role in driving withdrawal responses to von Frey filament/pinprick stimulation as well as tonic pain-indicating licking responses to skin pinch (Duan et al., 2014; Gatto et al., 2021). Surprisingly, both 75μl fCRD-evoked VMRs, measured via EMG recordings of abdominal muscle contractions, and 75μl fCRD-evoked aversion, measured via the step-down assay, were unchanged in VGLUT3^Lbx1^-Abl and SOM^Lbx1^-Abl mice compared with their respective control littermates (Figure S4).

Spinal CCK^Lbx1^ neurons were reported to drive withdrawal responses to air puffing, skin brushing, von Frey filament stimulation and pinprick applied to the hairy skin or to the hindpaw (Abraira et al., 2017; Gatto et al., 2021; Liu et al., 2018; Peirs et al., 2021). We first confirmed a marked deficit in withdrawal responses to von Frey filament stimulation in CCK^Lbx1^-Abl mice (Figure S5A), and then found marked attenuation in licking responses to skin pinch (Figure S5B), a stimulation producing tonic pain in humans (Huang et al., 2019). These phenotypes were not due to a general deficit in motor functions *per se*, since mutant mice and control littermates produced comparable withdrawal and licking responses in response to hot plate stimulation (Figure S5C and S5D). We then found that 75μl fCRD-evoked VMRs and aversion, observed in control littermates, were virtually abolished in CCK^Lbx1^-Abl mice (Figure S5E-H).

Collectively, these findings indicate a difference in spinal substrates in transmitting acute noxious mechanical information from the skin versus the rectum. Mechanical nociception and affective pain evoked from the skin are mediated via spinal neurons co-expressing SOM-Cre and CCK-Cre or via circuits connected by SOM and CCK lineage neurons, based on concurrent loss of these responses in SOM^Lbx1^-Abl and CCK^Lbx1^-Abl mice. In contrast, visceromotor reflexes and affective pain evoked by noxious rectal fCRD require spinal neurons expressing CCK-Cre, but not SOM-Cre or VGLUT3-Cre, based on their loss in CCK^Lbx1^-Abl mice and preservation in SOM^Lbx1^-Abl and VGLUT3^Lbx1^-Abl mice.

### Spinal VGLUT3 lineage neurons mediated low-intensity fCRD-evoked aversion in DSS-treated mice

We next investigated the roles of spinal VGLUT3 lineage in transmitting visceral information in mice with colitis. To help assess how spinal VGLUT3 lineage neurons responded to fCRD, we created *Vglut3*^*Cre*^-*tdTomato* mice by crossing *Vglut3*^*Cre*^ with a Cre-dependent tdTomato reporter. tdTomato^+^ neurons in the sacral spinal cord were enriched in the intermediate zone within the dorsal horn but also found in more deep laminae (Figure 2A). In naïve *Vglut3*^*Cre*^*-tdTomato* mice without DSS treatment, neither sham 0μl fCRD nor 25μl fCRD induced any FOS in tdTomato^+^ neurons (data not shown). In DSS-treated *Vglut3*^*Cre*^*-tdTomato* mice receiving sham 0μl fCRD, some FOS induction was observed in tdTomato^+^ neurons, and this FOS induction was significantly increased by 25μl fCRD (Figure 2A and 2B), observed at both lumbosacral (L6-S4) and coccygeal (Col1-3) levels (Figure S6). The total number of FOS^+^;tdTomato^+^ neurons from the lumbo-sacral-coccygeal spinal cord increased from 89.3 ± 3.5 with sham 0μl fCRD to 240.3 ± 15.6 with 25μl fCRD, a net increase of 151. If all 12 sets of spinal sections were analyzed, we estimated that 25μl fCRD activated around 1812 of VGLUT3^Cre^-tdTomato^+^ neurons in the lumbo-sacral-coccygeal spinal cord, although this number could be underestimated since FOS induction may not capture all neuronal activation.

**Figure 2.**
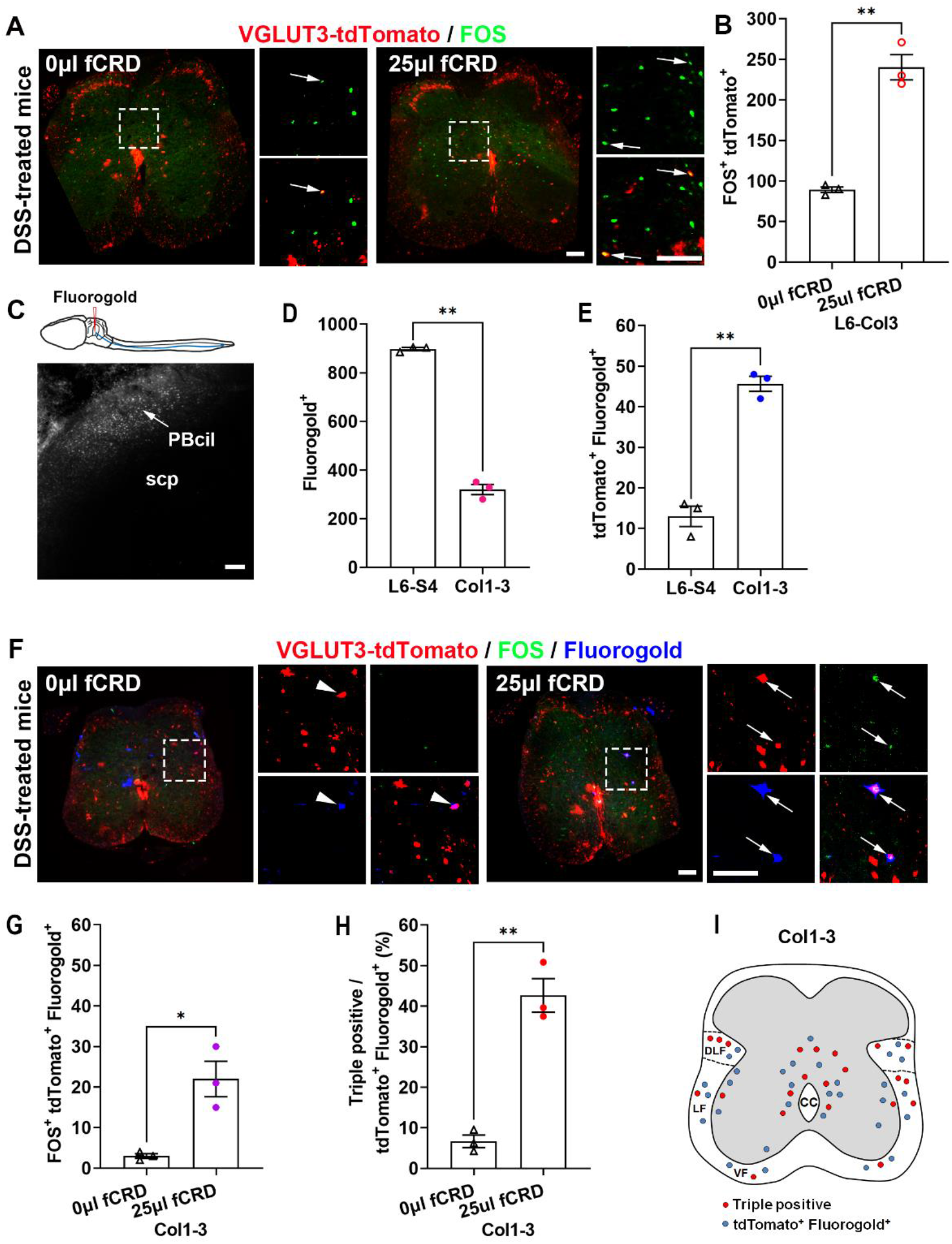
Responses of spinal VGLUT3 lineage neurons to low-intensity fCRD in DSS-treated mice. (A, B) Representative sections through the lumbo-sacral-coccygeal spinal cord of DSS-treated *Vglut3*^*Cre*^*-tdTomato* mice. Arrows indicating tdTomato^+^ neurons expressing FOS. Panel B shows an increase of FOS^+^;tdTomato^+^ neurons per set of sections through the lumbo-sacral-coccygeal spinal cord in response to 25μl fCRD compared with 0μl fCRD control (Student’s *t*-test, n = 3 per group, *t*_4_ = 9.45, **p < 0.01). (C-E) Retrograde labelling of spinal neurons projecting to the parabrachial nucleus (PBN). A representative section through the PBN of a *Vglut3*^*Cre*^*-tdTomato* mouse, showing Fluorogold injected into the central/internal-lateral PBN (“PBcil”). SCP: superior cerebellar peduncle. One of twelve sets of sections through the lumbo-sacral-coccygeal spinal cord were analyzed. PBcil-projecting neurons (Fluorogold^+^) were enriched in lumbosacral (“L6-S4”) compared with coccygeal (“Col1-3”) levels (D, Student’s *t*-test, n = 3 per group, *t*_4_ = 26.35, **p < 0.01), whereas PBcil-projecting VGLUT3 neurons (tdTomato^+^;Fluorogold^+^) were enriched in coccygeal compared with L6-S4 levels (E, Student’s *t*-test, n = 3 per group, *t*_4_ = 10.45, **p < 0.01). (F-H) Low-intensity fCRD induced FOS in PB-projecting VGLUT3 lineage neurons. Representative sections through the coccygeal cord from *Vglut3*^*Cre*^*-tdTomato* mice (F). Arrowhead and arrows indicating a FOS-negative and FOS-positive tdTomato^+^;Fluorogold^+^ neurons, respectively. 25μl fCRD caused an increase of FOS^+^;tdTomato^+^;Fluorogold^+^ neurons (G, Student’s *t*-test, n = 3 per group, *t*_4_ = 4.32, *p < 0.05) and an increase of the percentage of tdTomato^+^;Fluorogold^+^ cells with FOS (H, Student’s *t*-test, n = 3 per group, *t*_4_ = 8.15, **p < 0.01). (I) Schematic showing the distribution of PB-projecting VGLUT3 lineage neurons without FOS induction (blue dots, tdTomato^+^;Fluorogold^+^) or with FOS induction (red dots, FOS^+^;tdTomato^+^;Fluorogold^+^; “triple positive”) in the coccygeal cord. DLF: dorsal lateral funiculus. LF: lateral funiculus. VF: ventral funiculus. CC: central canal. Scale bars: 100μm. Data are shown as mean ± SEM.

Note that in response to 25μl fCRD, FOS induction was only observed in a very small subset of spinal VGLUT3 lineage neurons. We next asked if these activated neurons could contain ascending projection neurons, which are well known to represent a very small fraction of spinal neurons (Cameron et al., 2015; Polgar et al., 2010). As described later, spinal VGLUT3 lineage neurons do include projection neurons whose axons terminate in the central/internal-lateral subnuclei of the parabrachial nuclei (PBcil) (see below, Figure 4), a relay station crucial for affective pain transmission (Ma, 2022; Palmiter, 2018; Saper, 2000). To assess the location of PB-projecting spinal VGLUT3 lineage neurons and their responses to 25μl fCRD, we performed retrograde labeling by placing Fluorogold to the caudal PBcil region (Figure 2C), though the tracer also diffused to part of the cerebellum (data not shown). Two days later, these mice were fed with DSS for seven days and then subjected to 25μl fCRD or sham 0μl fCRD. From each set of spinal sections we analyzed, we detected 897.7 ± 6.4 and 320.0 ± 21.0 Fluorogold^+^ PBcil-projecting neurons at L6-S4 and coccygeal levels, respectively (Figure 2D), 90.3 ± 5.4 and 14.7 ± 1.5 of which resided in lamina I (data not shown). In contrast, for PBcil-projecting VGLUT3 neurons marked by tdTomato^+^;Fluorogold^+^, we detected 13.0 ± 2.5 and 45.7 ± 1.9 cells per set of L6-S4 and coccygeal sections, respectively (Figure 2E), showing enrichment in the coccygeal spinal cord and a notable exclusion from lamina I (see below, summarized in Figure 2I). If all twelve sets of spinal section analyzed, we estimate that there are around 156 and 548 PBcil-projecting spinal VGLUT3 lineage neurons within the lumbosacral and coccygeal segments of the spinal cord, respectively. We then found that following 25μl fCRD stimulation, among 45.7 ± 1.9 tdTomato^+^;Fluorogold^+^ neurons in the coccygeal spinal cord, 22.0 ± 4.4 of them showed FOS induction (Figure 2F and 2G). In other words, 42.7 ± 1.4% of PBcil-projecting VGLUT3 lineage neurons within the coccygeal cord responded to low-intensity fCRD in DSS-treated mice (Figure 2H), in comparison with 6.7 ± 1.5% in response to sham 0μl fCRD (Figure 2F-H). As summarized in Figure 2I, both FOS^+^ and FOS-negative PB-projecting spinal VGLUT3 lineage neurons reside in the dorsal lateral funiculus, the lateral funiculus, the ventral funiculus as well as the area surrounding the central canal.

**Figure 3.**
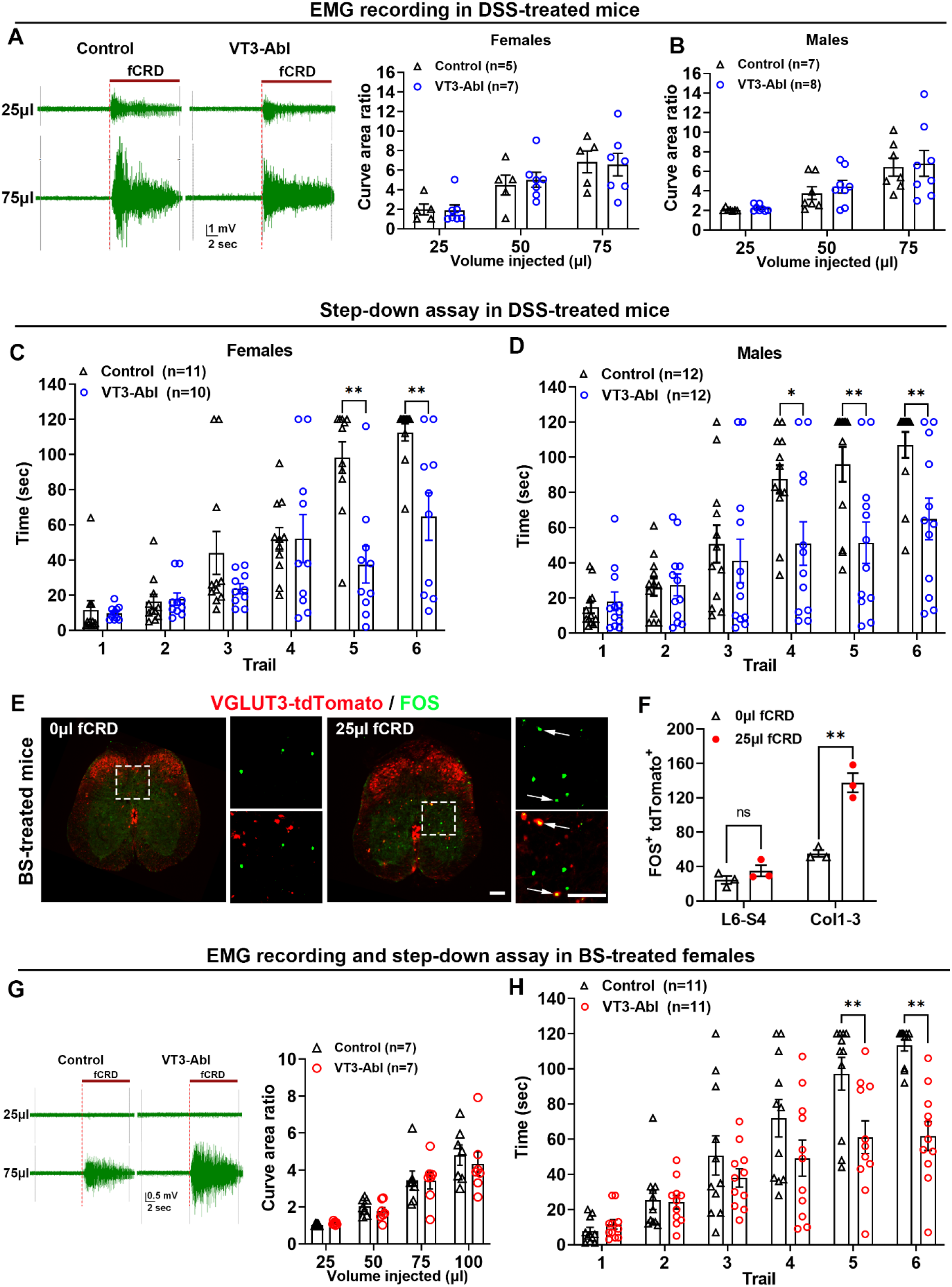
VGLUT3^Lbx1^ neurons drove aversion in mice with colitis or with central disinhibition, but not sensitized VMRs. (A, B) Ablation of VGLUT3^Lbx1^ neurons had no impact on VMRs. Representative traces of EMG activity evoked by 25 or 75 μl fCRD in DSS-treated females (A, left). No changes in evoked EMG activities in VGLUT3^Lbx1^-Abl (“VT3-Abl”) mice compared with control littermates, for both females (A, right, two-way ANOVA, n = 5, 7, *F*_2,20_ = 0.11, p = 0.89) and males (B, two-way ANOVA, n = 7, 8, *F*_2,26_ = 0.06, p=0.94). (C, D) No changes in 25μl fCRD-evoked aversive learning in DSS-treated VT3-Abl mice compared with control littermates, for both females (C) and males (D) (two-way ANOVA; females: n = 10, 11, *F*_5,95_ = 7.62, p < 0.01; males: n = 12 per group, *F*_5,110_ = 3.74, p < 0.01; post-hoc Holm-Sidak test: *p < 0.05, **p < 0.01). (E, F) 25μl fCRD-evoked FOS induction in spinal VGLUT3 neurons in mice receiving intrathecal injection of bicuculline and strychnine (BS). Representative coccygeal spinal sections from *Vglut3*^*Cre*^*-tdTomato* mice. Arrows indicating tdTomato^+^ neurons expressing FOS. Compared with sham distension, 25μl fCRD caused an increase of FOS^+^;tdTomato^+^ neurons (F, two-way ANOVA; n = 3 per group, *F*_1,4_ = 75.29, p = 0.001; post-hoc Holm-Sidak test: **p < 0.01; ns, not significant, p = 0.32). Scale bars: 100μm. (G) Representative traces of EMG activity induced by 25 or 75 μl fCRD in BS-treated females (G, left) and no changes in evoked EMG activity in VT3-Abl females versus control littermates (two-way ANOVA, n = 7 per group, *F*_3,36_ = 0.21, p = 0.89). (H) 25μl fCRD-evoked aversive learning following BS-induced central disinhibition was impaired in VT3-Abl females compared with control littermates (two-way ANOVA, n = 11 per group, *F*_5,100_ = 4.73, p < 0.01; post-hoc Holm-Sidak test: **p < 0.01). Data are shown as mean ± SEM.

**Figure 4.**
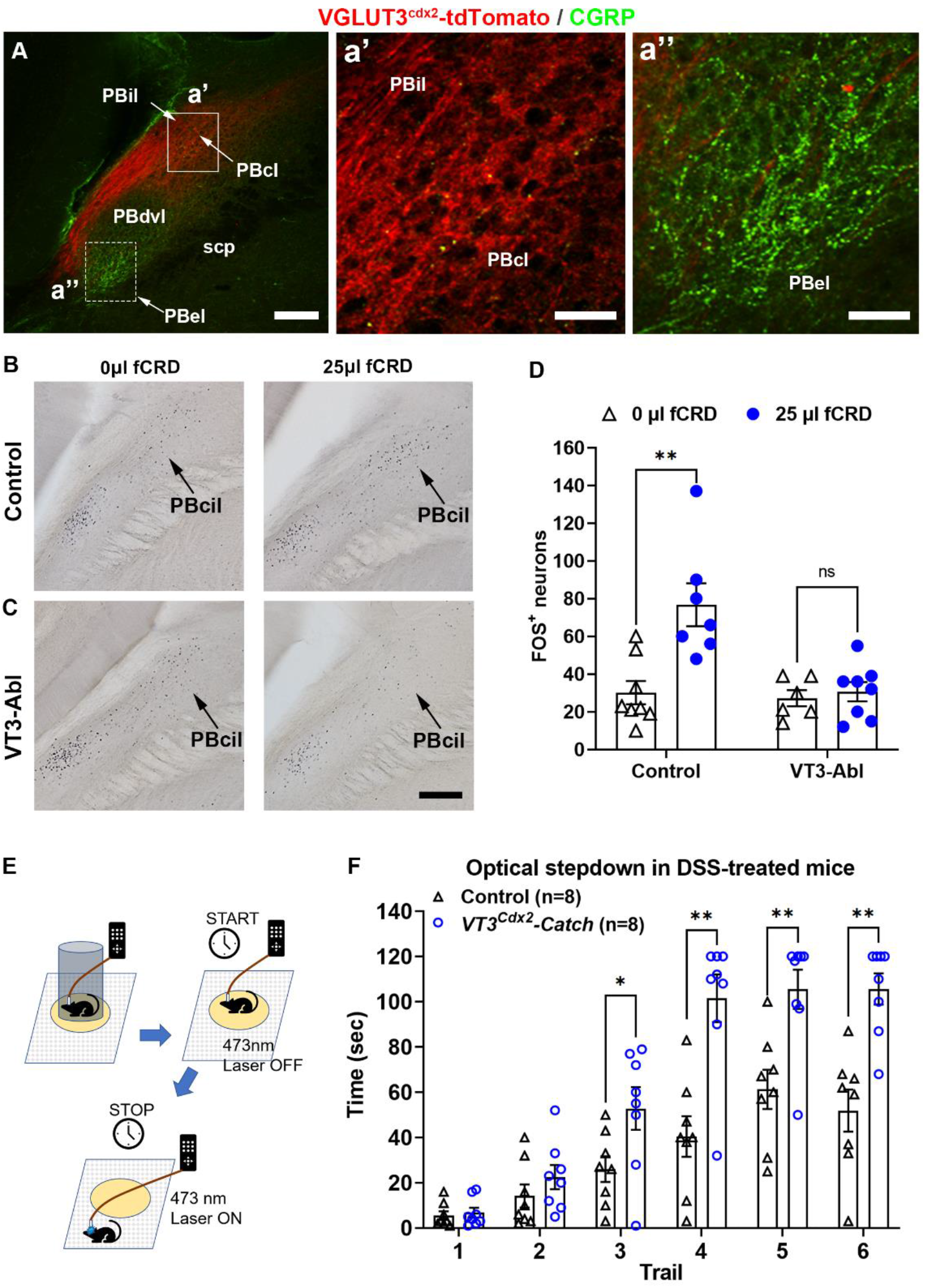
PB-projecting spinal VGLUT3 lineage neurons drove aversion. (A) A transverse section through the parabrachial nucleus of *Vglut3*^*Cdx2*^*-tdTomato* mice. Note dense tdTomato^+^ terminals (red) in the internal-lateral (PBil) and central-lateral (PBcl) subnuclei of the parabrachial nucleus (box a’), referred collectively to as PBcil, but not in part of the dorsal-ventral lateral PB nucleus (PBdvl) or the external-lateral parabrachial nucleus (PBel) marked by CGRP staining (box a’’, green). SCP: superior cerebellar peduncle. Scale bars: 100 μm in A, 20 μm in a’ and a’’. (B-D) Loss of 25μl fCRD-evoked FOS induction in the PBcil of DSS-treated VT3-Abl compared with control mice (two-way ANOVA, n = 6-8, *F*_1,14_ = 9.08, p < 0.01; post-hoc Holm-Sidak test: **p < 0.01; ns, not significant, p = 0.87). Scale bars: 100 μm. (E) Schematic description of the step-down aversive learning assay, with blue light delivery once a mouse stepped down from the petri dish (for details, see methods). (F) Optical stimulation of terminals in the PBcil drove aversive learning in DSS-treated *Vglut3*^*Cdx2*^*-CatCh* mice compared with control littermates (two-way ANOVA, n = 8 per group, *F*_5,70_ = 6.35, p < 0.01; post-hoc Holm-Sidak test: *p < 0.05, **p < 0.01). Data are shown as mean ± SEM.

We next examined the role of spinal VGLUT3 lineage neurons in transmitting visceral sensory information. EMG recordings showed that VMRs evoked by 25μl, 50μl, and 75μl fCRD were all unaffected in DSS-treated VGLUT3^Lbx1^-Abl mice compared with control littermates (Figure 3A and 3B). In stark contrast, 25μl fCRD-evoked aversive learning, measured via the step-down assay, was virtually abolished in DSS-treated VGLUT3^Lbx1^-Abl males and females (Figure 3C and 3D), with the residual learning curves comparable to that evoked by non-aversive 25μl fCRD in naïve mice (see above, Figure 1H and 1I). Other control experiments show that ablation of VGLUT3^Lbx1^ neurons did not affect colitis development, as indicated by similar body weight loss, comparable induction of myeloperoxidase (MPO, a biomarker for colitis), and unchanged locomotion activity (Figure S7A-F). Collectively, these studies show that spinal VGLUT3 lineage neurons are required to transmit rectal mechanical allodynia, but not sensitized visceromotor reflexes.

One hallmark of visceral pain is the manifestation of referred hypersensitivity in the cutaneous region (Cervero and Laird, 1999; Cui et al., 2022; Gebhart and Bielefeldt, 2016; Qiao and Tiwari, 2020). Consistently, DSS-treated control littermates showed cutaneous mechanical hypersensitivity, as indicated by reduced withdrawal thresholds in response to von Frey filament stimulation applied to the hindpaw (Figure S8A) and a gain of brush-evoked dynamic mechanical scores (Figure S8B). Such punctate and dynamic mechanical hypersensitivity was virtually abolished in DSS-treated VGLUT3^Lbx1^-Abl mice (Figure S8A and S8B). In other words, spinal VGLUT3^Lbx1^ neurons are required to transmit DSS-induced rectal mechanical allodynia as well as referred cutaneous mechanical hypersensitivity.

### Requirement of spinal VGLUT3 lineage neurons for the transmission of sensitized visceral pain induced by central disinhibition

Previous cutaneous pain studies showed that spinal VGLUT3 neurons are part of the polysynaptic pathways linking inputs from cutaneous low threshold mechanoreceptors to pain transmission neurons residing in superficial laminae, and such inputs are normally gated via feedforward activation of inhibitory neurons (Cheng et al., 2017; Peirs et al., 2021; Peirs et al., 2015). Most colorectal sensory neurons respond to low threshold mechanical stimulation, even if they encode sensory information at noxious ranges (Brierley et al., 2018; Christianson and Davis, 2010; Feng et al., 2010; Feng and Guo, 2020; Gebhart and Bielefeldt, 2016). To determine if analogous feedforward inhibition acted to prevent low-intensity fCRD from evoking pain, we analyzed mice with central disinhibition created via intrathecal injection of bicuculline and strychnine (BS) that blocked GABA_A_ and glycine receptors, respectively (Cheng et al., 2017). We found that in naïve wild type littermates, BS-induced disinhibition caused increased VMRs evoked by noxious 75μl and/or 100μl fCRD in comparison with mice without BS treatment (Figure S9A and S9B). 25μl CRD and 50μl fCRD were, however, still insufficient to produce sensitized VMRs (Figure S9A and S9B). In stark contrast, BS-induced central disinhibition did allow 25μl fCRD to produce sufficient negative teaching signal, as indicated by the step-down aversive learning assay (Figure S9C and S9D). Note that females appeared to display stronger aversion compared with males (Figure S9C and S9D), and we therefore focused on BS-treated females for subsequent studies.

We next investigated the roles of VGLUT3^Lbx1^ neurons in mediating BS-induced sensitized VMRs and aversion. We first found that in comparison with sham 0μl fCRD, 25μl fCRD was able to induce FOS in a subset of VGLUT3^Cre^-tdTomato^+^ neurons in BS-treated females and intriguingly, this induction occurred mainly in coccygeal levels, with minimal induction detected at L6-S4 levels (Figure 3E and 3F). We then found that sensitized VMRs evoked by noxious CRD (75μl and 100μl fCRD) were unaffected in BS-treated VGLUT3^Lbx1^-Abl females compared with control females (Figure 3G). In stark contrast, aversive learning evoked by low-intensity 25μl fCRD was virtually abolished in BS-treated VGLUT3^Lbx1^-Abl females (Figure 3H), with the residual learning curve (Figure 3H) comparable to that evoked by innocuous 25μl fCRD in naïve females (see Figure 1H). These data suggest that BS-sensitive feedforward inhibition provides a gate control under naïve conditions, preventing low-intensity rectal fCRD from driving VGLUT3^Lbx1^ neuron-dependent affective pain. Thus, under both inflammatory and central disinhibition conditions, spinal VGLUT3^Lbx1^ neurons are necessary for aversion evoked by low-intensity fCRD, whereas spinal neurons spared in VGLUT3^Lbx1^-Abl mice are sufficient to mediate sensitized visceromotor reflexes.

### Spinal VGLUT3^Lbx1^ neurons transmit fCRD-evoked affective information to the parabrachial nuclei

To understand how spinal VGLUT3^Lbx1^ neurons transmit affective visceral information, we first investigated if they include ascending projection neurons. To do this, we again used an intersectional genetic approach to create *Vglut3*^*Cdx2*^*-tdTomato* mice, in which tdTomato expression were confined to cells with coexpression of VGLUT3-Cre and Cdx2-Flpo (Figure S10A). Within the central nervous system, Cdx2-Flpo expression is confined to the spinal cord (Bourane et al., 2015), thereby allowing us to visualize central projections by spinal lineage neurons. It should be noted that for neuronal ablation studies, we used *Lbx1-Flpo* mice to generate VGLUT3^Lbx1^ mice, rather than using *Cdx2-Flpo* mice, since Cdx2-Flpo is also expressed in DRGs, whose use would lead to concurrent ablation of peripheral VGLUT3 lineage neurons in DRGs. Also, we did not create *Vglut3*^*Lbx1*^*-tdTomato* mice to study central projections, since Vglut3^Lbx1^ can label various trigeminal nuclei, including the paratrigeminal nuclei that locate right next to the parabrachial (PB) nuclei (Cheng et al., 2017), which might make it complicated to assess ascending projection exclusively from spinal VGLUT3 lineage neurons. We found that in *Vglut3*^*Cdx2*^*-tdTomato* mice, dense tdTomato^+^ terminals were observed in the central-lateral and internal-lateral parabrachial subnuclei (collectively referred to as PBcil) (Figure 4A), bypassing the exterior-lateral parabrachial subnuclei (PBel) defined by immunofluorescent staining of CGRP-the calcitonin gene-related peptide (Figure 4A) (Han et al., 2015) as well as an intermediate region that may correspond to part of the dorsal/ventral-lateral PB subnuclei (PBdvl) (Figure 4A). Innervation was also detected in the medial habenula nucleus (Figure S10B), but not in dorsal medial thalamic nuclei (Figure S10B), the ventral posterolateral thalamus (Figure S10C) or the periaqueductal gray (Figure S10D).

We then found that 25μl fCRD led to significant FOS induction in the caudal PBcil (Bregma: from -5.2 mm to -5.4 mm) in DSS-treated control littermates compared with sham 0μl fCRD (Figure 4B-D), but not in the more rostral PBN (Bregma: from -5.0mm to -5.2 mm) (data not shown). This FOS induction in the caudal PBcil was lost in DSS-treated VGLUT3^Lbx1^-Abl mice (Figure 4B-D), indicating that spinal VGLUT3^Lbx1^ neurons are required to transmit sensitized rectal sensory information to the caudal PBcil.

Neurons in the internal-lateral parabrachial nuclei (PBil, part of the a forementioned PBcil) have been strongly implicated in the transmission of the affective component of cutaneous pain (Barik et al., 2021; Deng et al., 2020; Huang et al., 2019; Ma, 2022). We next tested if spinal VGLUT3 neurons that project to the PBcil transmit an aversive signal. To test this, we generated *Vglut3*^*Cdx2*^*-Catch* mice, in which spinal VGLUT3 lineage neurons expressed the calcium translocating channelrhodopsin (CatCh) (Kleinlogel et al., 2011) plus the EYFP reporter (Figure S11A), and EYFP^+^ fibers can be observed in the PBcil area (Figure S11B). Again, *Cdx2-Flpo* mice, rather than *Lbx1-Flpo* mice, were used for optogenetic studies since the use of *Lbx1-Flpo* mice would drive CatCh expression in adjacent paratrigeminal nuclei. An optic fiber was surgically implanted on the mouse head with its tip targeting the caudal PBcil. Neurons in the lateral PB are known to be sensitized under chronic pain conditions (Chiang et al., 2019; Matsumoto et al., 1996; Raver et al., 2020; Uddin et al., 2018), and we accordingly focused on testing the impact of optical stimulation in DSS-treated *Vglut3*^*Cdx2*^*-CatCh* mice. We first found that 473nm blue light laser stimulation was able to induce Fos in the caudal PBcil in DSS-treated *Vglut3*^*Cdx2*^*-CatCh* mice compared with DSS-treated control siblings that lacked CatCh expression (Figure S11C and S11D), indicating that optical activation of these terminals was sufficient to activate PBcil neurons. We then conducted a modified step-down aversive learning assay, such that whenever a mouse stepped down from the petri dish to the floor, blue light was turned on to activate CatCh^+^ terminals in the caudal PBcil (Figure 4E). We found that this optical stimulation, at both 30 Hz (Figure 4F) and 15 Hz (Figure S11E), was sufficient to drive robust aversive learning in DSS-treated *Vglut3*^*Cdx2*^*-Catch* mice, requiring only 3-5 trials to produce conditioned avoidance, whereas the same optical stimulation produced minimal aversive learning in control littermates that lacked CatCh expression (Figure 4F). Without DSS treatment, optical stimulation produced a trend of aversive learning, almost reaching significance after six trials (Figure S11F), indicating the occurrence of sensitized information transmission from the VGLUT3^Cdx2^-CatCh^+^ terminals to the PBcil in mice with colitis, although sensitization could occur in more down downstream targets such as the ACC and the amygdala associated with aversive learning (Louwies et al., 2021; Lyubashina et al., 2022; Prusator and Greenwood-Van Meerveld, 2017; Thompson and Neugebauer, 2017, 2019; Wang et al., 2017; Yan et al., 2012). Importantly, this optical stimulation was insufficient to induce VMRs in DSS-treated *Vglut3*^*Cdx2*^*-Catch* mice (Figure S11G). Nor can it increase VMRs in response to 25-75μl fCRD in DSS-treated mice (Figure S11H). Thus, activation of PB-projecting spinal VGLUT3 lineage neurons can sufficiently produce aversion, without influencing visceromotor reflexes.

## DISCUSSION

Our studies suggest the existence of different spinal circuits that drive sensitized visceromotor responses versus affective visceral pain, echoing analogous circuit-level segregation in processing distinct dimensions of the cutaneous somatosensory system (Ma, 2022). Ablation of spinal VGLUT3 lineage neurons led to abolition of affective pain-indicating aversive learning in response to low-intensity rectal fCRD in mice with colitis or with spinal circuit disinhibition. In contrast, spinal neurons spared in VGLUT3^Lbx1^-Abl mice were sufficient to drive sensitized defensive visceromotor reflexes that might evolve to help get rid of harmful objects in the GI tract (Drake et al., 2015; Feng and Guo, 2020). Several studies shows that acute and chronic visceral inflammation can activate spinal neurons within lamina I that project to the nucleus of the solitary nucleus (NTS), marked by the expression of the neurokinin 1 receptor (NK1R) or the POU4F1 transcription factor (Gamboa-Esteves et al., 2004; Nishida et al., 2022; Zhang et al., 2013). These neurons could be attractive candidates for driving sensitized visceromotor responses since activation of the NTS is broadly involved with reflexive control of body physiology (Palmiter, 2018; Saper, 2000; Williams et al., 2016).

Spinal VGLUT3 lineage neurons include projection neurons whose axons terminate in the central/internal-lateral parabrachial subnuclei (PBcil), and low-intensity rectal fCRD in DSS-treated mice activates neurons in the caudal part of the PBcil. Interestingly, caudal PBcil-projecting VGLUT3 lineage neurons are enriched in the coccygeal spinal cord, 43% of which responded to low-intensity rectal fCRD following colitis induction. Photoactivation of these terminals was sufficient to produce aversion in DSS-treated mice. Neurons in the internal-lateral parabrachial nuclei (part of PBcil) send direct projections to the dorsomedial thalamic complex (DMC) (Barik et al., 2021; Bester et al., 1999; Bourgeais et al., 2001; Deng et al., 2020; Fulwiler and Saper, 1984; Huang et al., 2019; Ma, 2022), which in turn send projections to ACC (Ma, 2022; Price, 2002; Xiao and Zhang, 2018). In humans, lesions of the medial thalamus or the ACC caused pain indifference, losing strong unpleasantness and emotional distress in response to injury-causing stimuli (Ballantine et al., 1967; Foltz and White, 1962; Freeman and Watts, 1948; Mark et al., 1960; Mark et al., 1963; Young et al., 1995). We speculate that fCRD-evoked mechanical allodynia might partially process through the PB-DMC-ACC pathways. Consistently, lesions in the ACC mimicked ablation of spinal VGLUT3 lineage neurons, leading to a selective loss of CRD-evoked aversive learning in animals with colitis, without affecting visceromotor reflexes (Yan et al., 2012). In other words, measurement of visceromotor reflexive responses alone would fail to reveal a crucial role of spinal VGLUT3 neurons and ACC neurons in affective visceral mechanical allodynia. Our studies, however, do not argue against the measurement of defensive visceromotor reflexes *per se*. For the cutaneous somatosensory system, circuits normally associated with acute reflexive-defensive responses can in fact gain ability to access affective pain pathways under pathological conditions and contribute to the development of comorbidities, such as fear and anxiety (Ma, 2022). Rather, our studies suggest that multiple physiological and behavioral assays are needed to study neural substrates processing nociceptive and affective dimensions of visceral sensory information as well as pain comorbidities.

Spinal VGLUT3 lineage neurons may represent a convergent node for driving mechanical allodynia originating from different tissues. A subset of these neurons that reside in deep laminae form polysynaptic pathways that link inputs from cutaneous Aβ low threshold mechanoreceptors (Aβ-LTMRs) to pain transmission (T) neurons residing in superficial laminae, and such inputs are normally gated via feedforward inhibition; following nerve lesions, Aβ-LTMR inputs gain the ability to activate VGLUT3 neurons and drive one of most prevalent and bothersome forms of neuropathic pain: the morphine-resistant dynamic allodynia evoked by tactile stimulation across a skin area (Cheng et al., 2017; Peirs et al., 2021; Peirs et al., 2015). Notably, recent single cell gene expression profiling shows that many colorectal DRG neurons express Piezo2 (Hockley et al., 2019; Meerschaert et al., 2020), a low threshold mechanically gated ion channel (Jiang et al., 2021; Kefauver et al., 2020). Upon blockage of GABA and glycine receptors, low-intensity fCRD gained ability to produce aversion in a VGLUT3 neuron-dependent manner, suggesting that inputs from low-threshold colorectal afferents to spinal VGLUT3 neurons are also gated via feedforward inhibition. Future studies will be directed to determine if DSS-induced inflammation might cause a reduction of inhibitory inputs and/or an increase of excitatory inputs to spinal VGLUT3 lineage neurons, such that low-threshold colorectal distension can sufficiently activate VGLUT3 neurons, followed by T neuron activation to drive affective pain. One hallmark for inflammatory visceral pain manifestation is the development of referred cutaneous mechanical hypersensitivity (Bai et al., 2019; Bielefeldt et al., 2006; Bourdu et al., 2005; Cao et al., 2021; Gao et al., 2021; Jain et al., 2022; Laird et al., 2000; Laird et al., 2002; Lv et al., 2019; Traub et al., 2008; Zhou et al., 2008). Such referred hypersensitivity was also abolished in VGLUT3^Lbx1^-Abl mice, suggesting that VGLUT3 neurons must be part of previously proposed spinal circuits receiving convergent inputs from cutaneous and visceral afferents (Cervero, 2009; Cui et al., 2022; Fang et al., 2021; Fuentes and Christianson, 2018; Gebhart and Bielefeldt, 2016; Luz et al., 2015; Qiao and Tiwari, 2020). Thus, spinal VGLUT3 neurons represent an attractive cellular target for treating allodynia originated from both cutaneous and rectal tissues.

Our studies also gain new insight into spinal substrates transmitting acute noxious mechanical information evoked from the skin versus the rectal regions. Pinprick and pinch, which produce sharp and tonic pain from the skin, respectively (Ma, 2022), fail to produce pain from the colon and rectum in humans (Feng and Guo, 2020; Gebhart and Bielefeldt, 2016; Lewis, 1942). Not coincidently, spinal SOM lineage neurons, which are necessary for skin pinprick-evoked withdrawal responses and skin pinch-evoked tonic pain-indicative licking responses (Duan et al., 2014), are dispensable for visceromotor reflexes and aversion evoked by noxious rectal fCRD. Consistently, SOM neurons locate mainly in lamina II (Duan et al., 2014), a region with rare innervation from visceral afferents (Cervero and Connell, 1984). For spinal lamina I neurons with convergent inputs from cutaneous and colorectal mechano-nociceptors, this convergence might be due to more direct inputs from visceral nociceptors and indirect inputs from cutaneous mechanical nociceptors through lamina II SOM neurons. The identities and roles of those lamina I neurons with monosynaptic inputs from somatic and visceral afferents (Luz et al., 2015) need further investigation. In contrast, spinal CCK lineage neurons must be more heterogenous and as a group, they are required to transmit noxious mechanical information from both the skin and the rectum (Figure S5) (Abraira et al., 2017; Gatto et al., 2021; Liu et al., 2018; Peirs et al., 2021). Future studies will be directed to identify the SOM-negative subset of CCK lineage neurons for crucial transmission of noxious visceral mechanical information.

### Limitations of the study

The crucial roles of spinal VGLUT3 lineage neurons in driving affective visceral mechanical allodynia are suggested collectively from a set of observations: i) a virtual loss of low-intensity fCRD-evoked aversive learning in VGLUT3^Lbx1^-Abl mice, ii) a projection of these neurons to the central/internal-lateral parabrachial nuclei (PBcil) that have been implicated in affective pain processing, iii) a high response rate of these PB-projecting neurons to low-intensity fCRD in mice with colitis, iv) a loss of fCRD-evoked FOS induction in the caudal PBcil in VGLUT3^Lbx1^-Abl mice, and v) the sufficiency of VGLUT3 terminal activation in the PBcil for driving aversion in mice with colitis. Despite these strengths, our studies do contain limitations. Firstly, VMRs were measured under anesthetic conditions, and VMRs can be impacted differentially with different anesthetic compounds (Ness and Gebhart, 1988). VMRs need to be tested in wake animals in the future, although restraint stress associated with wake animal studies could create its own complexity, capable of modulating both VMRs and pain (Cao et al., 2021; Johnson et al., 2020). Secondly, more studies are needed to determine if spinal VGLUT3 neurons could mediate mechanical allodynia evoked from the colon, not just from the rectum. Recent gene expression profiling studies have revealed the presence of i) shared DRG neuron subtypes innervating proximal and distal segments of colon and rectum and ii) unique subtypes innervating preferentially in the rectum segment (Hockley et al., 2019; Meerschaert et al., 2020). At this moment, it is still unclear if the affective visceral allodynia evoked by low-intensity fCRD in the rectum is mediated by unique and/or shared visceral afferent subtypes. Nonetheless, the virtual loss of referred mechanical hypersensitivity in VGLUT3^Lbx1^-Abl mice suggests that these neurons might be sensitized, or be connected to spinal neurons sensitized, by visceral afferent inputs from the GI tract beyond the rectum. Thirdly, our studies focused on mechanical allodynia evoked by low-intensity fCRD in mice with colitis. Other assays need to be used or developed to measure spontaneous pain as well as comorbidities, such as anxiety, depression and cognitive deficits (Gebhart and Bielefeldt, 2016; Johnson et al., 2020; Wang et al., 2017; Yuan and Greenwood-Van Meerveld, 2021). Fourthly, we have not yet ruled out a contribution from vagal afferents to low-intensity fCRD-evoked aversion in mice with colitis. Our fCDR was performed in distal rectum with minimal innervation from vagal afferents. This method does allow noxious fCRD to evoke VMRs and aversion in a vagal nerve-independent manner (but see Figure S2A on the complexity in interpreting results from vagotomy). For allodynia evoked by low-intensity rectal fCRD in DSS-treated mice, the situation could be more complex. On one hand, spinal VGLUT3 lineage neurons, necessary for driving sensitized aversion, do contain ascending projection neurons terminating in the internal-lateral parabrachial nuclei that do not receive inputs from the nucleus of the solitary tract (NTS), the relay station for vagal afferents (Saper, 2000). On the other hand, VGLUT3 lineage neurons might include interneurons connected to ascending projection neurons that terminate in brain nuclei receiving inputs from vagal afferents, such as the external lateral parabrachial nuclei and the amygdala that involve with aversive learning as well (Campos et al., 2018; Chiang et al., 2019; Chiang et al., 2020; Kang et al., 2022; Palmiter, 2018; Saper, 2000). As such, inputs from vagal afferents from the GI tract beyond the rectum could contribute to sensitization in those convergent brain centers that in turn enables spinal afferents to drive sensitized aversive learning. Future studies are needed to test this hypothesis. Finally, our studies have not yet examined if rectal fCRD causes mechanical tension in surrounding somatic tissues, such as fascia that wraps the external anal sphincter muscle and attaches these muscles to tailbones. In other words, rectal fCRD might in fact represent a model to study anorectal pain that contains mixed visceral and somatic components, whose management remains a significant medical challenge (Knowles and Cohen, 2022; Lee et al., 2017). Despite these unsolved issues, our studies identify VGLUT3 lineage neurons as the crucial spinal substrate for the transmission of affective inflammatory pain evoked by distension starting at the rectal region, irrespective of containing a somatic component or not.

## SUPPLEMENTARY INFORMATION

Supplemental information includes 11 figures and can be found with this article online at

## ACKNOWLEDGMENTS

We thank Bradford B. Lowell, Martyn Goulding, Susan M. Dymecki, and the Allen Brain Institute/the Jackson Laboratory for genetically modified mice. The work was supported primarily by the NIH grant (DK122833) and partially by the Wellcome Trust grant (200183/Z/15/Z) to Q.M.

## AUTHOR CONTRIBUTION

L.Q. and S.L. designed and performed all experiments. Q.M. conceptualized and supervised the study. S.L., L.Q. and Q.M. wrote the manuscript.

## STAR* METHODS

### KEY RESOURCE TABLE

#### RESOURCE AVAILABILITY

##### Lead Contact

Further information and requests for resources and reagents should be directed to and will be fulfilled by the Lead Contact, Qiufu Ma.

##### Materials Availability

- The study did not generate new plasmids or unique reagents
- The study did not generate new mouse lines; the mouse lines used for intersectional manipulations were all acquired from public resources or from relevant investigators (see below, “Mice” in “EXPERIMENTAL MODEL AND SUBJECT DETAILS”).

##### Data and Code Availability

This study did not generate any unique datasets or code.

### EXPERIMENTAL MODEL AND SUBJECT DETAILS

### Mice

Animal experiments were performed with protocols approved by the Institutional Animal Care and Use Committee at Dana Farber-Farber Cancer Institute and followed NIH guidelines. Mice were kept with a 12-h/12-h light/dark cycle and had ad libitum access to standard laboratory mouse pellet food and water, and in a temperature- and humidity-controlled room. The *Vglut3-IRES-Cre* (referred to as *Vglut3-Cre*) mice were acquired from Dr. Bradford Lowell (Cheng et al., 2017). The *SOM-IRES-Cre* (referred here to as *SOM-Cre*; JAX #013044), *CCK-IRES-Cre* (referred here to as *CCK-Cre*; JAX#012706), *R26-CAG*^*LSL-tdTomato*^ (“Ai14”, JAX#007908), *R26*^*ds-tdTomato*^ (“Ai65”, JAX#021875) and *R26*^*ds-CatCh*^ reporter mice (“Ai80”, JAX#025109) were acquired from the Jackson Laboratory. The *Lbx1*^*Flpo*^, *Cdx2*^*Flpo*^, *Tau*^*ds-DTR*^ mice were acquired from Dr. Martyn Goulding (Bourane et al., 2015; Britz et al., 2015). To ablate spinal *Vglut3 or SOM* lineage neurons, 8-10-week-old triple heterozygous mice (*Vglut3*^*Cre/+*^;*Lbx1*^*Flpo/+*^*;Tau*^*ds-DTR/+*^ or *SOM*^*Cre/+*^;*Lbx1*^*Flpo/+*^*;Tau*^*ds-DTR/+*^) were injected intraperitoneally with DTX (50 μg/kg, dissolved in 200μl saline, Sigma-Aldrich, D0564) twice with a 72-hour interval. To ablate *CCK* lineage neurons and due to DTR expression in muscles, 8-10-week-old triple heterozygous mice (*CCK*^*Cre/+*^;*Lbx1*^*Flpo/+*^*;Tau*^*ds-DTR/+*^) were injected intrathecally with DTX (5ng in 20μl saline, Sigma-Aldrich, D0564) twice with a 72-hour interval.

Control littermates without DTR expression received same DTX injections. All behavioral assays, electromyography recording, and histochemical analyses were performed at least 28 days after the second injection. For genetic ablation or optogenetic studies, littermates lacking either the *Flpo* or the *Cre* allele were used as sibling controls. The ablation efficiency and specificity of spinal VGLUT3 and SOM lineage neurons using this intersectional strategy had been verified previously (Cheng et al., 2017; Duan et al., 2014). All these mice resulted from the intersectional genetic crossing were of mixed genetic backgrounds, containing 129 and C57BL6. Wide type mice used to generate data in Figure 1 also came from the intersectional genetic breeding involved with *Vglut3-Cre*. Males and females were analyzed separately for most studies, with the exception that i) for optogenetic stimulation of central terminals in the parabrachial nuclei, males and females showed similar effects and the data were combined, and ii) for aversive learning following central disinhibition induced by bicuculline and strychnine, females were chosen after pilot studies revealed apparently weaker aversive learning by males. No estrus cycle was evaluated for females. For all EMG and behavioral analyses, investigators were blinded to the genotypes and experimental conditions.

### METHOD DETAILS

#### Retrograde tracing from proximal colon and distal rectum

To trace DRG neurons and vagal neurons residing in the nodose ganglia that innervated the colon or the rectum, 2% Fluorogold (FG, Fluorochrome) was injected into the muscle wall (5 spots, 0.5 μl per spot), using a glass micropipette pulled from the borosilicate glass (BF150-86-10; Sutter Instrument, Novato, CA, USA) with a Sutter P-1000 pipette puller. Mice were anesthetized with intraperitoneal injection of the ketamine/xylazine cocktail (87.5/12.5 mg per kg of body weight), and the lower abdominal cavity was opened with a 1-cm midline incision. For the colon, injections were performed at the backside of the colon, adjacent to mesenteric attachment, or at the frontside, opposite to mesenteric attachment, with the entry point 3.5 cm away from the anus. For the rectum, backside or frontside injections were again performed, using different batches of mice, with the entry point 0.5 cm away from the anus. For both injections, the pipette tip was pushed rostrally for about 0.4 cm into the muscle. All injections were performed under the microscope. Five days later, mice were sacrificed with CO_2_ and perfused with 50 ml PBS and then 50 ml 4% PFA. Nodose ganglia and DRGs (from thoracic T10 to sacral S3) were dissected and cryoprotected in 30% sucrose overnight at 4°C. Whole-mount fluorescent images of collected ganglia were taken.

#### Retrograde tracing from the parabrachial nuclei

Adult *Vglut3*^*Cre*^*-tdTomato* mice were anesthetized with intraperitoneal injection of the ketamine/xylazine mixture (87.5/12.5 mg per kg of body weight) mixture. Each mouse was then placed on a stereotaxic apparatus (Stoelting), with body temperature maintained at 37°C via a heating pad. A glass micropipette, pulled from borosilicate glass (BF150-86-10; Sutter Instrument, Novato, CA, USA) with a Sutter P-1000 pipette puller, was used to inject Fluorogold (400 nl for each side, 2% in water; Fluorochrome) bilaterally into the caudal part of the central/internal-lateral parabrachial nuclei, using the coordinate (AP: -5.3mm; ML: -1.3mm; DV: -2.3mm). After surgery, the analgesic compound (Meloxicam, 5 mg/kg) was given every 24 hours for two times. After a two-day recovery, mice were subjected to DSS-feeding for 7 consecutive days (see below) and then used for examining FOS induction in response to low-intensity fCRD (see below).

#### Recording fCRD-induced EMG responses

Electromyographic (EMG) recordings of abdominal muscle contractions were carried out under the urethane-induced mild sedation condition (intraperitoneal injection, 1.3g/kg of body weight), using a modified focal colorectal distension method (Annahazi et al., 2012). To deliver fCRD, a latex balloon made of a condom rubber patch (with the outer balloon diameter to be ∼1.7 mm before distention) was attached to the tip of the PE-50 tubing (ID = 0.58 mm, OD = 0.97 mm). A steel cannula (with 0.5 mm diameter, cut from 21G syringe needle) was inserted inside the tip of the tubing to provide a force-resistant base such that the silk thread can be tied tightly. The other end of the tubing was connected to a 100μl Hamilton micro-syringe filled with water. Special attention was paid to avoid air bubble in the balloon, the connecting tubing and the syringe. The balloon was inserted into the rectal position 0.5 cm away from the annus and the tubing was taped to the tail base. After urethane sedation induction and hair shaving, a 1-cm incision was performed to expose the left abdominal muscle. EMG activity in response to balloon distension was recorded with two electrodes inserted into the external oblique muscle. fCRD stimulations were applied by injecting water into the balloon with progressively increasing volume (25, 50, 75, 100μl) with at least 1-min interval between trials. Each fCRD was performed when baseline EMG activity was stable. Ten-second periods of EMG activity were recorded, right before and during balloon distension, and the curve area ratio was determined as the ratio of the spike area from these two recorded periods. All animals were euthanized after EMG measurements. For EMG data analyses, mice with high background activity, with spike amplitude higher than 1mV, which occurred in a small subset of mice, were excluded, and the exclusion was performed before genotypes or experimental groups were decoded.

#### The fCRD-induced step-down assay

A modified step-down aversive learning assay (Ness and Gebhart, 1988) was used to measure the affective component of visceral pain. Mice were habituated for three consecutive days to reduce stress associated with rectal balloon insertion. With light anesthesia by 2% isoflurane, the latex balloon was inserted into the rectum (0.5 cm away from the anus) with the connecting tube tapped to tail base, and the mouse was placed onto a limited open space for 10-min habituation (on the top of an upside-down 2-litter beaker to prevent the mouse from turning its head to bite the tube). On the step-down training day, each mouse first went through 10-min habituation process. The mouse was then put onto an elevated circular platform (an inverted peri dish with 1-cm height and 15-cm diameter) and capped with a plastic cylinder (14 cm in diameter x 16 cm height). The platform was positioned at the center of a square mesh (40 × 40 cm). Once the cap was lifted, the time (in seconds) mouse took to step down from the platform with four limbs to the mesh was recorded (the cut-off time is 120 seconds). Upon stepping down, the mouse immediately received fCRD delivered through the balloon with a defined water volume for 30 seconds and was then placed back to the habituation area after fCRD was stopped by drawing water back to the syringe. After a 5-min interval, the mouse went through the next trial, and in total, 6 consecutive trials were carried out. A progressive increase of time each mouse staying onto the platform was used as a measurement of avoidance learning.

#### DSS feeding

Mice were fed with 3% dextran sodium sulfate (DSS) in drinking water for 7 days to induce ulcerative colitis. Body condition score (BCS) and body weight (BW) were monitored daily. DSS was replaced with water on Day 8 and behavioral tests such as open field locomotion (see below) or the step-down assay (see above) were carried out on Day 9. For the step-down assay, 3-day habituation started on day 6 of DSS feeding. On Day 10, the mice were subjected to the measurement of fCRD-evoked EMG responses and then euthanized.

#### Immunohistochemistry

To characterize FOS induction by fCRD in the spinal cord and/or in the parabrachial nuclei (PBN), mice underwent the same habituation process for 3 days, as described in the step-down assay (see above). fCRD with fixed intensity was then delivered for 30 seconds per trial and 6 trials in total were performed with 5-min intervals. One and half hours later, mice were euthanized by CO_2_ and perfused with 50 ml of ice-cold PBS and then 50 ml of 4% paraformaldehyde (PFA) prepared in PBS, pH7.4. The brain and the lumbo-sacral-coccygeal spinal cord were dissected and fixed in 4% PFA for overnight at 4°C and then cryoprotected for another overnight in 30% sucrose in PBS. Tissues were then embedded in OCT (Thermo Scientific, 6502).

For the ABC-DAB-Nickel staining of the PBN, the brain was sectioned in a vibratome (Leica VT1000S) at 75 μm thickness and collected in PBS. Free-floating brain sessions were first bleached for 30 minutes with 0.3% H_2_O_2_ in PBS and then washed with PBST (PBS containing 0.1% Triton X-100) for 3 times. Sections were then blocked in PBST containing 5% goat serum at room temperature for 60 min and then incubated with the primary antibody diluted in blocking solution overnight at 4°C. Brain slices were then washed 3 times with PBST and incubated for 1 hour at room temperature with the biotinylated goat anti-rabbit secondary antibody. After three PBST washes, samples were incubated in the Avidin-Biotin pre-mix solution (VECTASTAIN Elite ABC Kits, Vector Labs, #PK-6101; 1:1000 in PBS) for 1 hour. Sections were then washed with PBS for three times, and dark blue signals were visualized in the Nickel-DAB solution (8mg NH_4_Cl, 20mg β-D-glucose, 1mg glucose oxidase in 10ml PBS containing 0.05% DAB and 0.05% nickel ammonium sulfate). For each mouse brain, all sections through the PBN region were analyzed, and FOS^+^ neurons in the rostral and caudal halves of the superior/central/internal-lateral subnuclei were counted.

For immunofluorescence staining, the lumbo-sacral-coccygeal spinal cord (from L6 to the end) was sectioned into twelve adjacent sets of sections at 30 μm thickness, one of which (containing 40-42 sections) was used for FOS staining. Air dried sections on a slide were permeabilized with PBST, incubated for 1h at room temperature with blocking solution (PBST containing 5% goat serum) and then incubated overnight at 4°C with primary antibodies in blocking solution. Primary antibody staining was detected and visualized with fluorophore–conjugated secondary antibodies, and these immunostaining signals plus the tdTomato and/or Fluorogold signals were captured using the Zeiss AX10 fluorescent microscope. For immunofluorescence staining on sections through the PBN (anti-CGRP in *Vglut3*^*Cdx2*^*-tdTomato* mice, and anti-FOS or anti-GFP in *Vglut3*^*Cdx2*^*-Catch* mice), the brain region covering the PBN was sectioned into six adjacent sets of sections at 25 or 30 μm thickness, and one of these six sets was analyzed. Representative sections through the caudal PBN were selected for visualizing anti-CGRP signals (plus tdTomato) or anti-GFP signals, as well as for quantifying FOS induction in the PBcil region in response to optical stimulation (see below), with FOS^+^ neurons per section counted and presented.

Primary antibodies used in present study: rabbit anti-FOS (Millipore, ABE457, 1:1000 for immunofluorescent and 1:5000 for DAB staining), mouse anti-CGRP (Sigma, C7113, 1:1000) and rabbit anti-GFP (Invitrogen, A11122, 1:1000 for immunofluorescent). Secondary antibodies: donkey anti-rabbit IgG-Alexa 488 (Jackson ImmunoResearch, 711-545-152, 1:500), donkey anti-mouse IgG-Alexa 488 (Jackson ImmunoResearch, 715-546-151, 1:500), donkey anti-rabbit IgG-Alexa 594 (Jackson ImmunoResearch, 711-585-152, 1:500) and biotinylated goat-anti-rabbit IgG (ABC kit, Vector Labs, BA-1000, 1:1000).

#### Intrathecal injection of bicuculline and strychnine

Mice were first anesthetized by 2% isoflurane and hairs on the caudal back were shaved. After identifying the midline of the lumbar enlargement, 31G needle connected to a 25 μl Hamilton microsyringe was punched into the subdural space, with successful puncture indicated by tail flicking response. 10 μl saline containing 0.02 μg of bicuculline (Sigma-Aldrich, 14340) and 0.05 μg of strychnine (Sigma-Aldrich, S0532) was injected, and isoflurane exposure were then stopped immediately. Ten minutes later, 25μl fCRD-evoked step-down assay or 25μl fCRD-evoked EMG activity in the external oblique muscle was recorded (see above), using different batches of mice.

#### Stereotaxic surgery for optic fiber implantation

For stereotaxic surgery, each mouse was anesthetized with intraperitoneal injection of the ketamine/xylazine cocktail (87.5mg/12.5 mg per kg of body weight). Mouse head was mounted on the stereotaxic frame (Stoelting), and custom-made optical fiber (material: Ceram; ferrule O.D: 2.5 mm; fiber core: 400μm; fiber length: 3.0mm; ThorLabs) was implanted unilaterally above the caudal central/internal-lateral parabrachial nucleus (AP: -5.3mm; ML: -1.3mm; DV: -2.0mm from brain surface). After 10 days of recovery, fiber-implanted mice were subjected to histochemical studies (see above) or the optical step-down assay (see below).

#### Optical step-down learning

473-nm blue light (30 Hz or 15 Hz, 20-ms pulse width, square wave, 10 mW; Opto Engine, Laser Model PSU-III-LED) was delivered through the optic fiber in *Vglut3*^*Cdx2*^*-Catch* mice. The step-down apparatus is the same as the one used for the fCRD-evoked step-down assay, except that the cap was replaced with an open roof and fCRD stimulation was replaced with blue light laser stimulation for 30 seconds.

#### Open field locomotion

Open field locomotion was recorded in an open arena (40 × 40 cm) with a camera in the central top. Without any habituation, each mouse was placed into the open field and video-recorded for 15 minutes. The horizontal distance travelled (in meters) was analyzed by Any-Maze software. Data were presented with 3-minute blocks.

#### ELISA to measure MPO

After 7-day DSS feeding, the distal rectum (0-1cm from annus) was dissected, weighted, and frozen immediately in dry ice. Tissue proteins were extracted by the lysis buffer containing 0.02M tris-hydrochloric acid (pH=8.0), 0.15M sodium chloride and 0.01% Triton X-100. After homogenization and centrifugation, the supernatant was collected and analyzed with the mouse myeloperoxidase ELISA kit (Thermo Scientific, EMMPO). Five pairs of DSS- and water-treated male and female mice were analyzed, and each sample was analyzed in triplicate and the average amount of MPO (ng per mg of tissue) was presented.

#### Measurement of cutaneous mechanical and thermal sensitivity

Withdrawal thresholds to von Frey filament stimulation, withdrawal latency or the duration of licking to 50°C hot plate and licking duration to skin pinching by CCK^Lbx1^-Abl mice and their control littermates were measured as described previously (Huang et al., 2019). For referred mechanical sensitivity measurement, mice were habituated for 30 minutes in a restricting cube (6 × 6 × 6 cm) on a wire mesh platform one day before DSS feeding. Baseline punctate and dynamic mechanical sensitivity was then measured as previously described (Cheng et al., 2017). After baseline measurement, mice were fed with 3% DSS for 7 days. Two days after completion of DSS feeding, punctate and dynamic mechanical hypersensitivity were measured.

### QUANTIFICATION AND STATISTICAL ANALYSIS

#### Statistical analysis

Results were expressed as mean ± SEM. Statistical analyses were performed by using the SigmaStat 3.5 and GraphPad Prism 9 software. All datasets were tested for normality and equal variance. For two-sample tests, data were analyzed with two-side Student’s unpaired *t*-test. For one-factor multiple comparisons, one-way ANOVA was used. For two-factor experiments in fCRD-induced EMG responses and fCRD-induced step-down learning, two-way ANOVA with repeated measurements was used, and if main effect reached significant, post hoc Holm-Sidak test was followed for pairwise comparisons. For the description of the correlation between balloon sizes and injected water volumes, the Pearson correlation test was used. The *P* < 0.05 was accepted as statistically difference.

