## supplemental figure for "Spinal VGLUT3 lineage neurons drive visceral mechanical allodynia but not visceromotor reflexes"

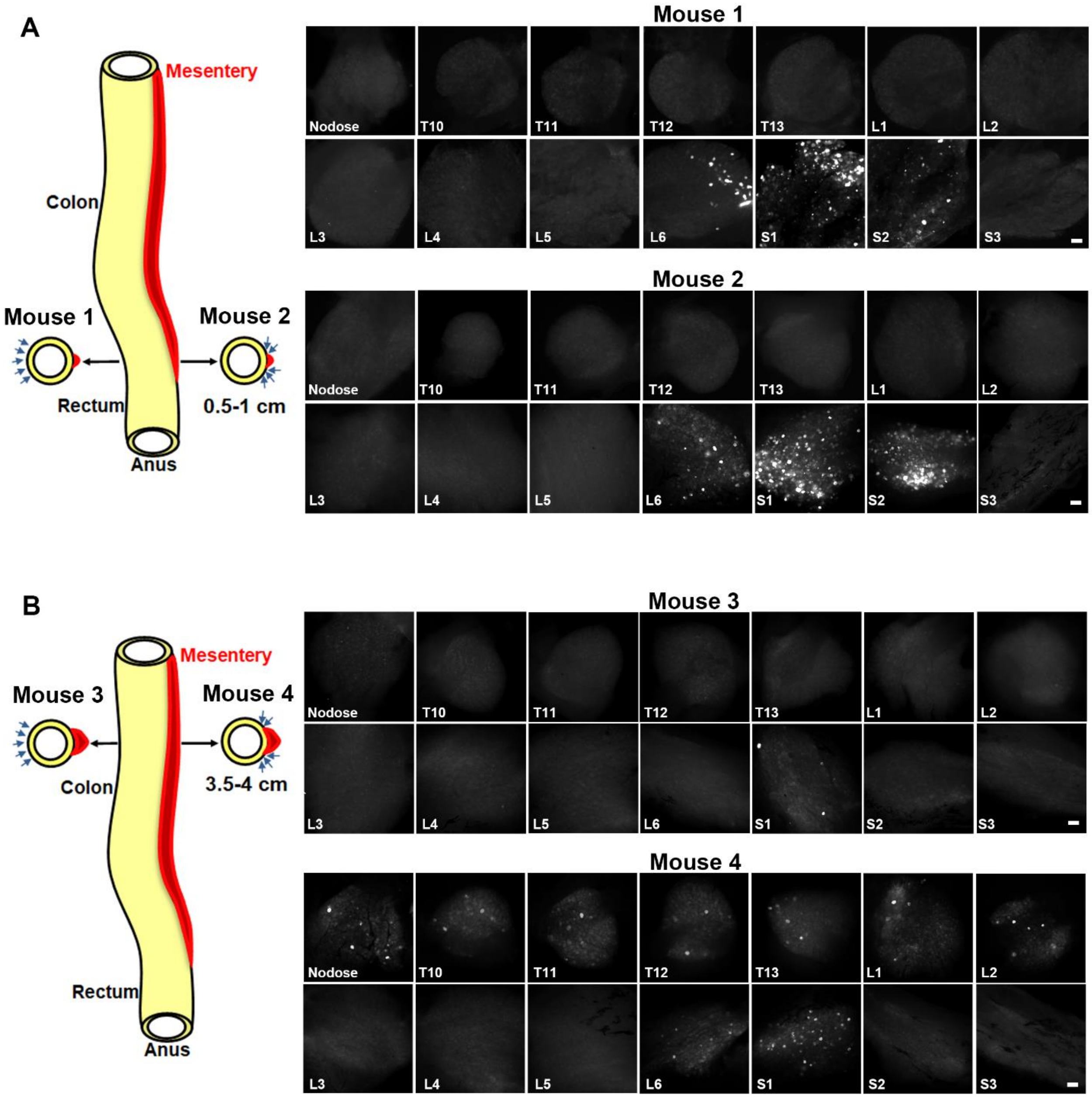


**Figure S1. Retrograde labeling of neurons in nodose ganglia and DRGs from colorectal regions. Related to Figure 1.**

(A) Distribution of retrogradely labeled cells following Fluorogold injections in the distal rectum (0.5-1 cm away from the anus), at either frontside or backside. Left, schematics showing injection spots. Both injections led to labeling in lumbosacral (L6-S2) DRGs, but none in nodose ganglia or thoracolumbar (T10-L5)/sacral (S3) DRGs. This side-independent innervation pattern is consistent with previous reports that rectal afferents innervate throughout the circumferential axis of distal rectum (Brierley et al., 2018; Feng and Guo, 2020). Our data are also consistent with progressive reduction of vagal afferent innervation along the proximal-distal axis of the gastrointestinal (GI) tract (Wang and Powley, 2000) and predominant innervation in the distal rectum by lumbosacral DRGs (Kyloh et al., 2011). However, other investigators reported more extensive labeling in nodose ganglia and in T10-L2 DRGs following tracer injection 1 cm away from the anus (Christianson et al., 2006 ; Meerschaert et al., 2020; Robinson et al., 2004). The nature behind this controversy remains unclear, but it might be due to a difference in tracer volumes, in terms of the total volume injected (2.5 μl in our studies versus 5-10 μl for studies by Christianson et al. and Meerschaert et al.) and the volume per injected spot (0.5 μl for each of 5 spots for our studies and 2-3 or 5 μl per spot by Christianson et al. and Meerschaert et al.). Potentially, large volume injection would be easier to cause tracer leakiness. Future studies are needed to clarify this controversy.

(B) Distribution of retrogradely labeled neurons following Fluorogold injections to the frontside or backside of the proximal colon (3.5-4 cm away from the anus). Left, schematic showing injection spots. Note that backside injection led to retrograde labeling in nodose ganglia, T10-L2 DRGs and L6-S1 DRGs, whereas frontside injection only produced minimal labeling in the S1 DRG. The enriched innervation within the backside of colon is consistent with previous electrophysiological recordings (Brierley et al., 2018; Feng and Guo, 2020). Scale bars: 100 μm.

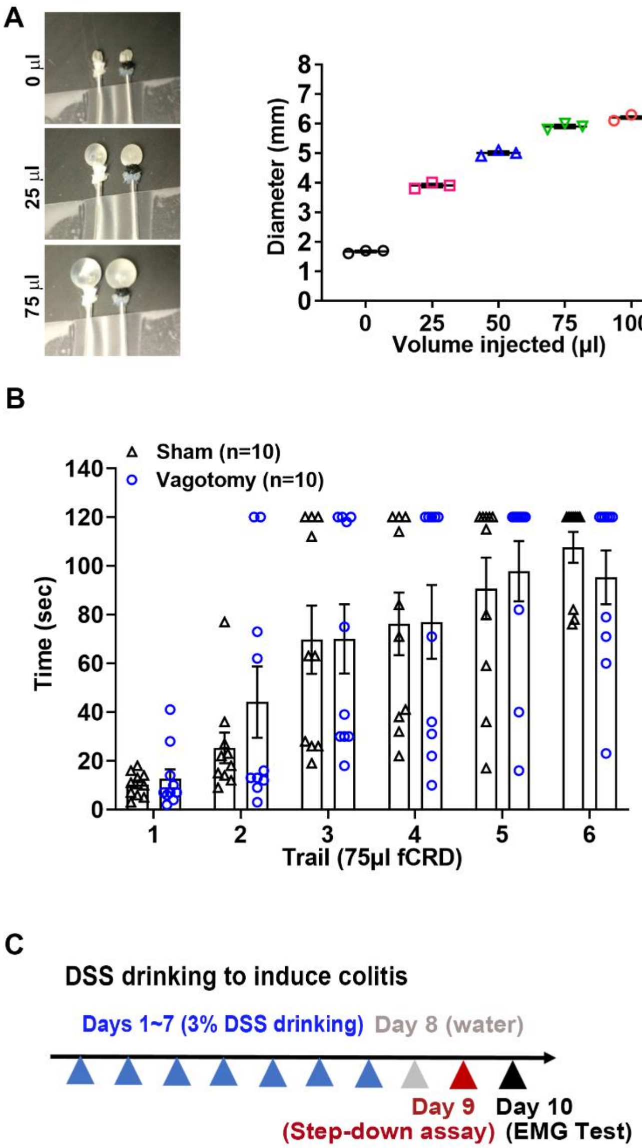


**Figure S2**. **The fCRD method, vagal nerve-independent aversion evoked by noxious fCRD and schematic description of DSS-induced colitis. Related to Figure 1.**

(A) The fCRD method is modified from Annahazi et al. (Annahazi et al., 2012). Left pictures showing distended balloons following water injection with volumes at 0, 25 and 75 µl. Three batches of homemade balloons show correlation coefficient between balloon diameters and injected water volumes (the Pearson correlation test, $r = 0.95$, $p < 0.05$). Note high reproducible distension from batch to batch.

(B) No impact of vagotomy on aversive learning evoked by 75 µl fCRD compared with intact mice using the step-down assay (two-way ANOVA, n = 10 per group, $F_{5,90} = 0.45$, $p = 0.81$).

This vagal nerve-independent aversion appears to be consistent with minimal vagal afferent innervation in the rectum (see Figure S1), although it is not known if this noxious CRD can cause mechanical tension in more proximal colonic regions and activate vagal afferents innervating there. Interpretation of vagotomy needs caution as well. Vagal afferents are functionally heterogeneous, with some afferents driving aversion (Cao et al., 2012; Palmiter, 2018) and other afferents playing an anti-nociception role, as indicated by sensitized nocifensive responses in mice with vagotomy, including CRD-evoked visceromotor responses (Chen et al., 2008; Furuta et al., 2009; Gschossmann et al., 2002). Thus, alternative explanations for the lack of vagotomy on fCRD-evoked aversion could be due to functional redundancy between spinal and vagal afferents and/or due to a cancellation effect by concurrent transection of distinct vagal afferents with opposing roles.

(C) Schematic illustration of colitis induction by DSS feeding, and the timing for the subsequent step-down test and EMG recording (for details, see methods). Data are shown as mean ± SEM.

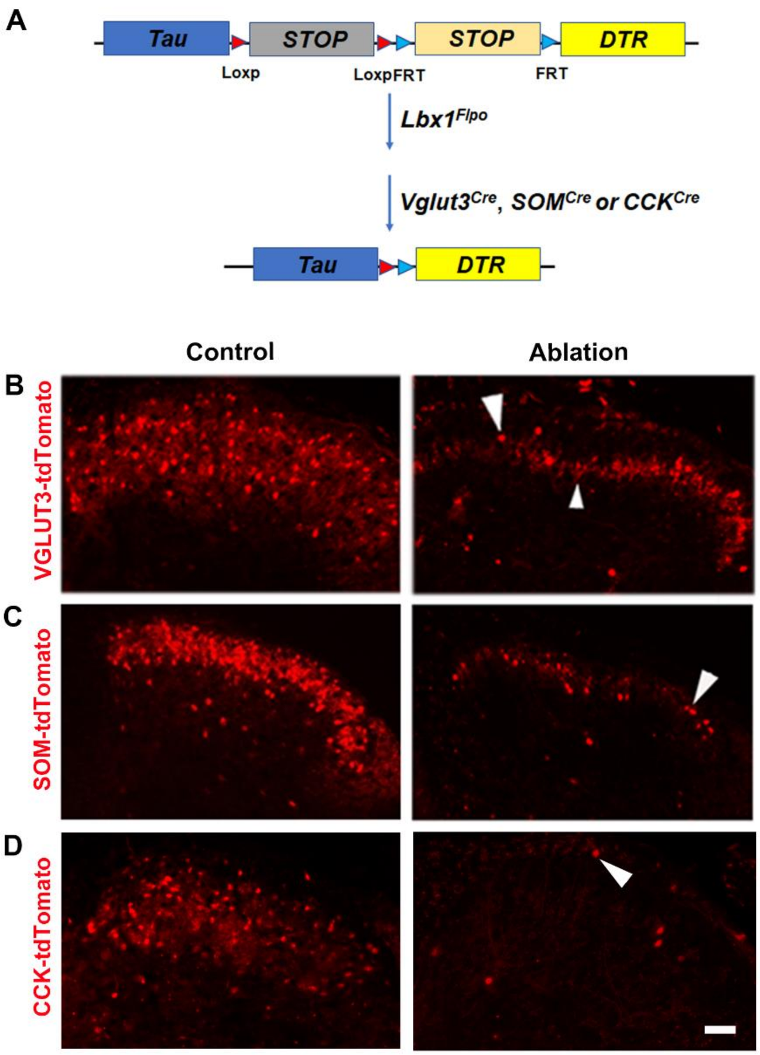


**Figure S3. Generation of VGLUT3$^{Lbx1}$-Abl, SOM$^{Lbx1}$-Abl and CCK$^{Lbx1}$-Abl mice. Related to Figure 1.**

(A) Schematic illustration of the intersectional genetic approach to target the human diphtheria toxin receptor (DTR) in the spinal neurons defined by co-expression of Lbx1-Flpo plus VGLUT3-Cre, SOM-Cre or CCK-Cre, following removal of two STOP cassettes via Cre- and Flpo-mediated DNA recombination, referred to as *Vglut3$^{Lbx1}$-DTR*, *SOM$^{Lbx1}$-DTR* or *CCK$^{Lbx1}$-DTR* mice.

(B, C, D) Not shown is the presence of a Cre-dependent tdTomato allele to label all Cre-expressing cells with tdTomato, for helping monitor cell ablation. After 2 rounds of intraperitoneal or intrathecal injection of the diphtheria toxin in *Vglut3$^{Lbx1}$-DTR, Som$^{Lbx1}$-DTR* or

*CCK^{Lbx1}-DTR* mice, the majority of VGLUT3-tdTomato$^{+}$, SOM-tdTomato$^{+}$, or CCK-tdTomato$^{+}$ neurons in the dorsal spinal cord were ablated. Large arrowhead in B and arrowheads in C and D indicate few remaining neurons, and small arrowhead in B indicating the central nerve terminals derived from unablated VGLUT3-tdTomato$^{+}$ DRG neurons. Scale bars: 100 μm.

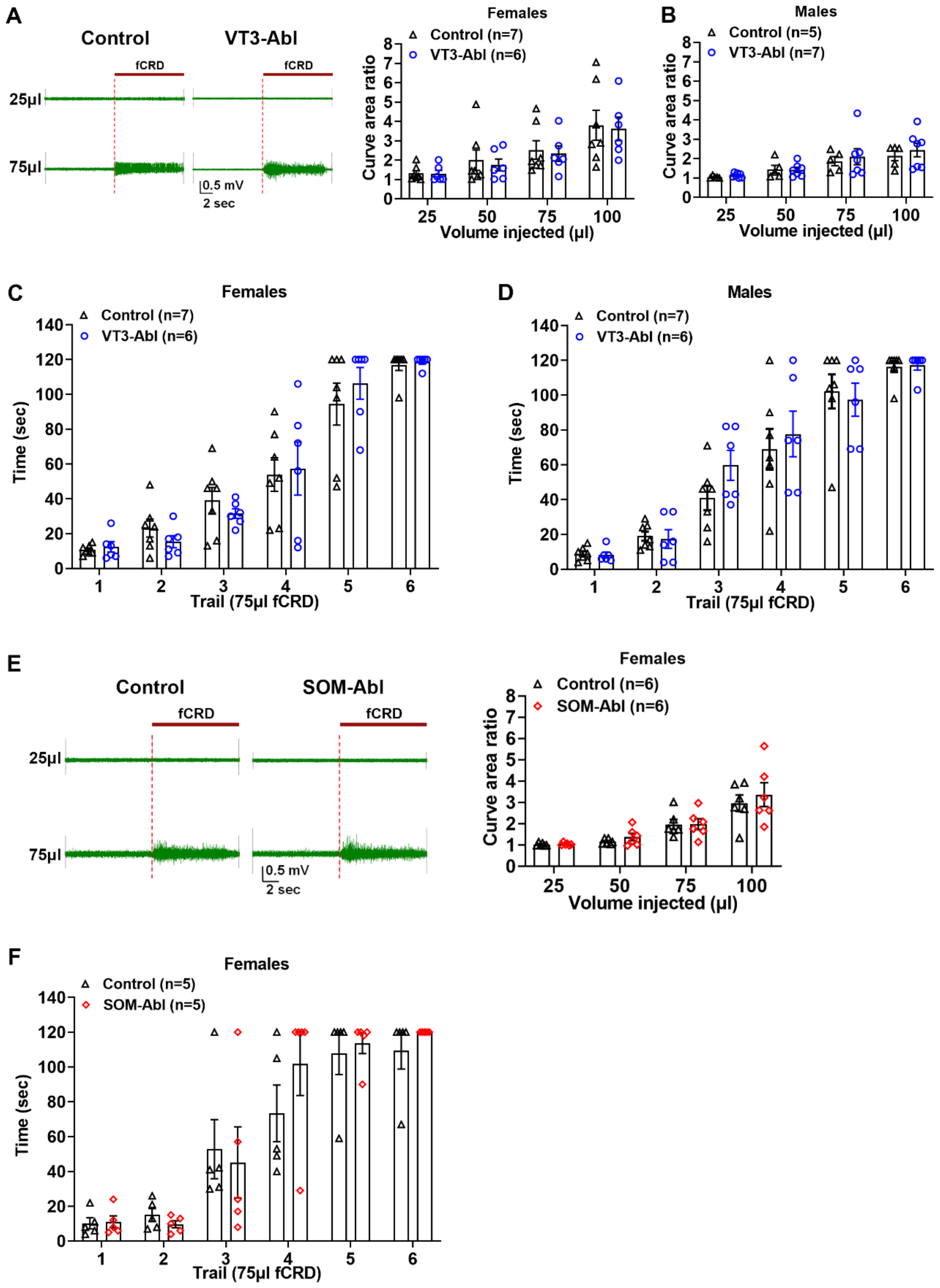


**Figure S4**. **Spinal VGLUT3$^{Lbx1}$ and SOM$^{Lbx1}$ neurons were dispensable for acute VMRs and aversion evoked by noxious 75µl fCRD. Related to Figure 1.**

(A, B) Ablation of VGLUT3$^{Lbx1}$ neurons had no impact on fCRD-evoked acute VMRs. Representative traces of EMG activity induced by 25 or 75 μl fCRD in naïve and VGLUT3$^{Lbx1}$-Abl ("VT3-Abl") females (A, left). No changes in EMG activities in VT3-Abl mice compared with control littermates, for both females (two-way ANOVA, females: n= 6, 7, $F_{3,33}$ = 0.04, p = 0.99;) and males (two-way ANOVA, males: n = 5, 7, $F_{3,30}$ = 0.35, p = 0.79).

(C, D) Compared with control littermates, no changes in 75μl fCRD-evoked aversive learning (the step-down assay) in both VT3-Abl females and males (two-way ANOVA; females: n = 6, 7, $F_{5,55}$ = 0.52, p = 0.76; males: n = 6, 7, $F_{5,55}$ = 0.90, p = 0.49).

(E, F) Ablation of SOM$^{Lbx1}$ neurons did not affect fCRD-evoked acute VMRs and aversion. Representative traces of EMG activity induced by 25 or 75 μl fCRD in naive and SOM$^{Lbx1}$-Abl ("SOM-Abl") females (E, left). Compared with control littermates, SOM-Abl females showed no changes in evoked EMG activities (E, right, two-way ANOVA, n= 6 per group, $F_{3,30}$ = 0.32, p = 0.81) or 75μl fCRD-evoked aversive learning via the step-down assay (F, two-way ANOVA, n = 5 per group, $F_{5,40}$ = 0.69, p = 0.63).

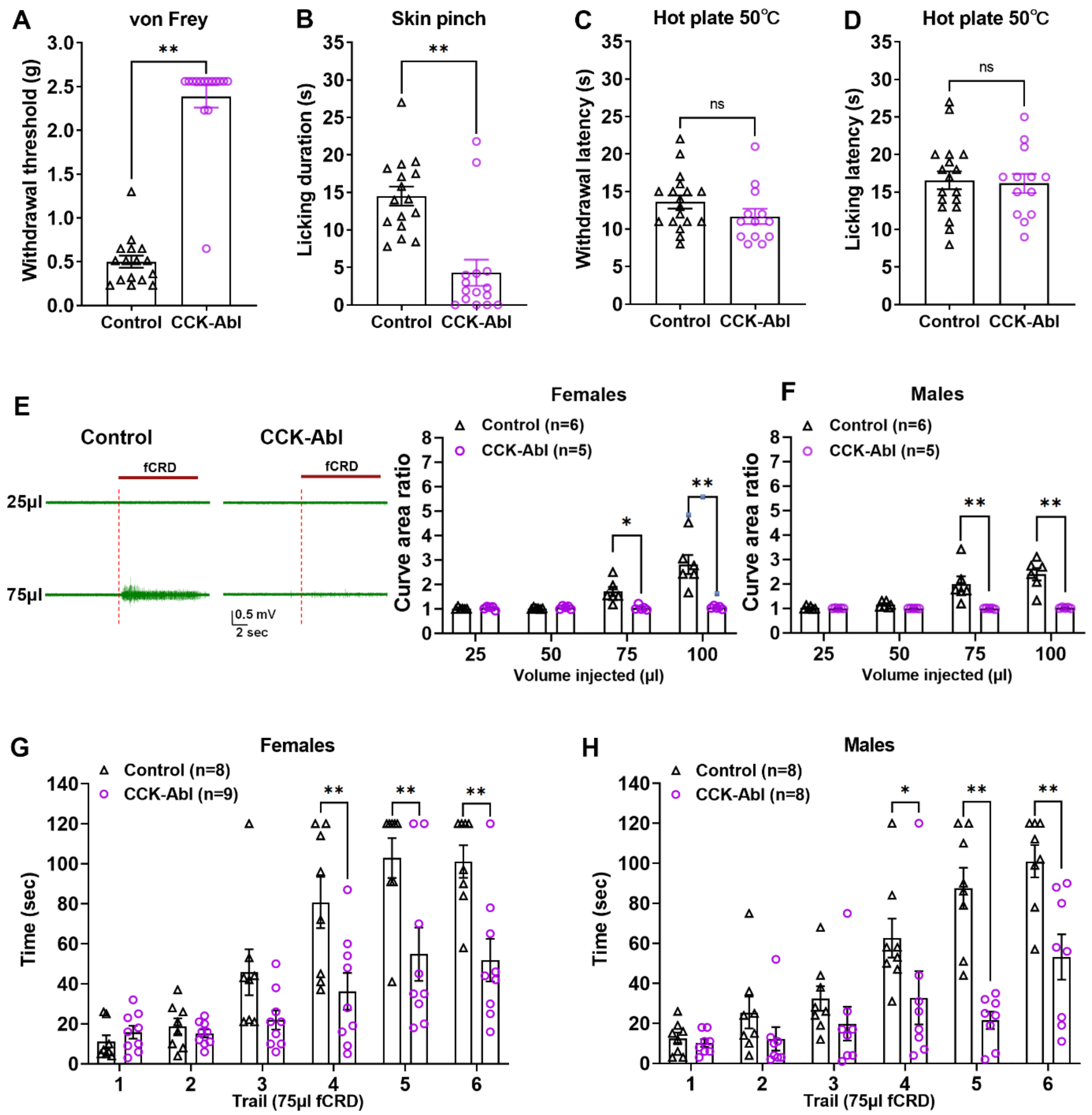


**Figure S5. Spinal CCK$^{Lbx1}$ neurons transmitted noxious mechanical information from both the skin and the rectum. Related to Figure 1.**

(A, B) CCK$^{Lbx1}$-Abl mice ("CCK-Abl") showed marked increase in withdrawal thresholds in response to von Frey filament stimulation applied to the hindpaw (Student's *t*-test, n = 15, 16, $t_{29}$ = 13.26, **p < 0.01) and a marked reduction in licking responses evoked by inescapable skin pinch (Student's *t*-test, n = 15, 16, $t_{29}$ = 4.80, **p < 0.01).

(C, D) CCK-Abl mice produced comparable withdrawal responses (Student's *t*-test, n = 13, 18, $t_{29} = 1.42$, p = 0.17) and licking responses (Student's *t*-test, n = 13, 18, $t_{29} = 0.23$, p = 0.82) in exposure to the 50℃ hot plate.

(E, F) CCK-Abl mice showed marked reduction in VMRs evoked by 75μl and 100μl fCRD. Representative traces of EMG activity induced by 25 or 75 μl fCRD in naive and CCK-Abl females compared with control females (E, left). No difference was detected for both females (E) and males (F) (two-way ANOVA, females: n= 5, 6, $F_{3,27} = 11.47$, p < 0.01; males: n= 5, 6, $F_{3,27} = 7.89$, p < 0.01; post-hoc Holm-Sidak test: *p < 0.05, **p < 0.01).

(G, H) Compared with control littermates, 75μl fCRD-evoked aversion was attenuated in both $CCK^{Lbx1}$-Abl females and males (two-way ANOVA; females: n = 8, 9, $F_{5,75} = 4.44$, p < 0.01; males: n = 8 per group, $F_{5,70} = 4.58$, p < 0.01; post-hoc Holm-Sidak test: *p < 0.05, **p < 0.01).

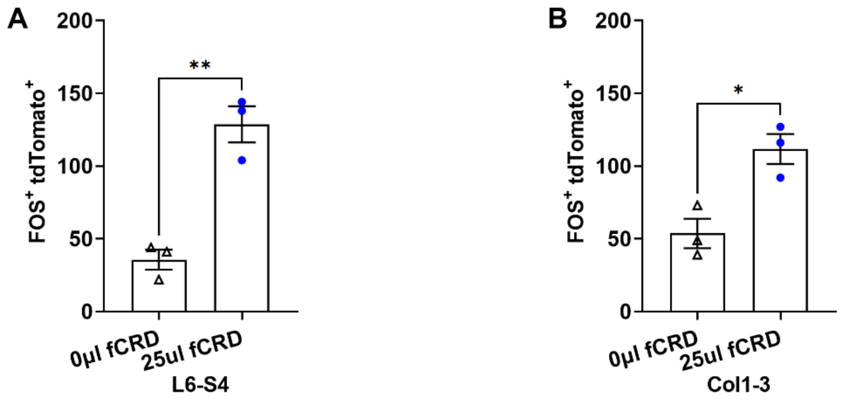


**Figure S6. Low-intensity fCRD-evoked FOS induction in VGLUT3 lineage neurons at both lumbosacral and coccygeal levels in DSS-treated mice. Related to Figure 2.**

(A, B) Compared with sham 0µl fCRD, 25µl fCRD in DSS-treated *Vglut3$^{Cre}$-tdTomato* mice led to an increase of FOS$^{+}$;tdTomato$^{+}$ neurons at both lumbosacral ("L6-S4") levels (Student's *t*-test, n = 3 per group, $t_4 = 6.53$, **$p < 0.01$) and coccygeal ("Col1-3") levels (Student's *t*-test, n = 3 per group, $t_4 = 4.02$, *$p < 0.05$).

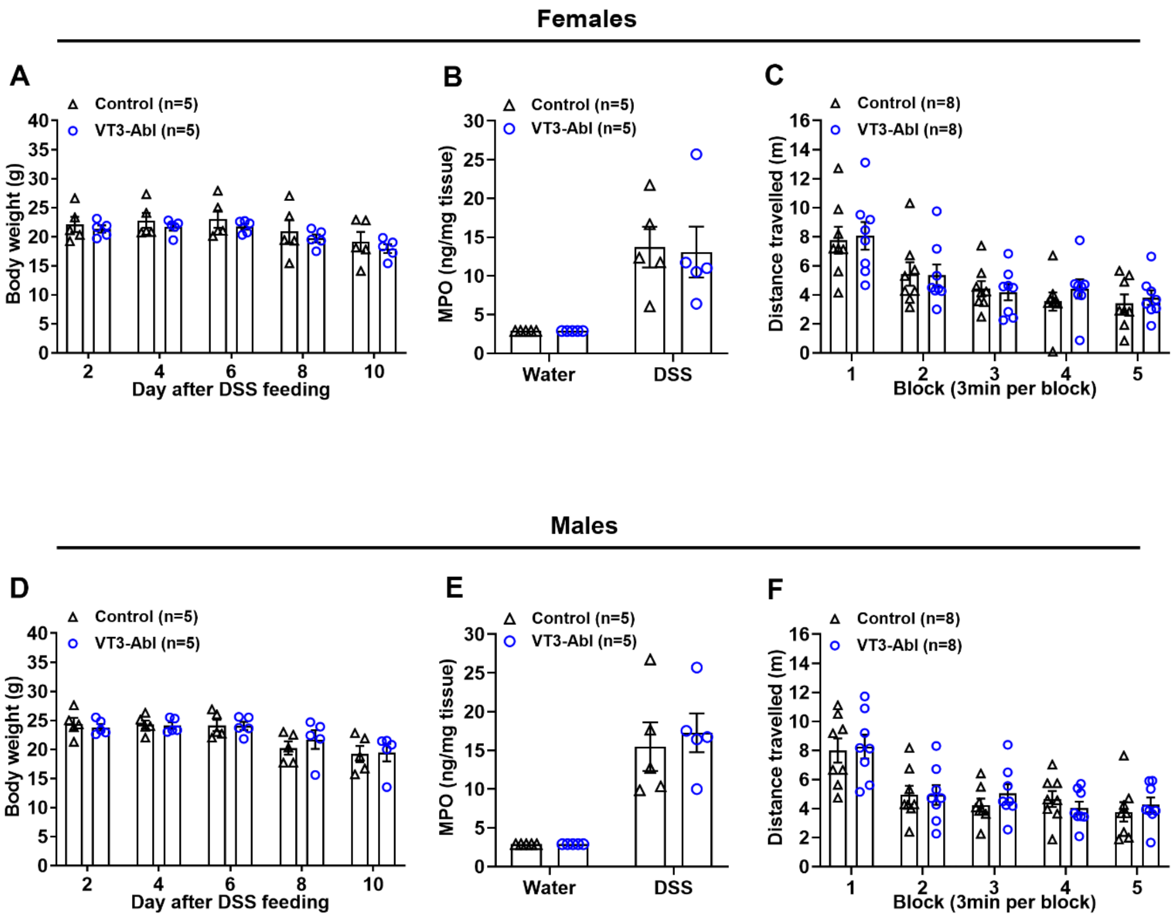


**Figure S7. Ablation of spinal VGLUT3 lineage did not affect DSS-induced colitis development. Related to Figure 3.**

(A-F) Following DSS feeding, $VGLUT3^{Lbx1}$-Abl ("VT3-Abl") mice and control littermates showed no difference in body weight (A, D; two-way ANOVA; females: n = 5 per group, $F_{4,32} = 0.08$, p = 0.99; males: n = 5 per group, $F_{4,32} = 0.70$, p = 0.60), MPO induction (B, E; two-way ANOVA; females: n = 5 per group, $F_{1,8} = 0.02$, p = 0.88; males: n = 5 per group, $F_{1,8} = 0.20$, p = 0.67), or locomotion activity (C, F; two-way ANOVA, females: n = 8 per group, $F_{4,56} = 0.22$, p = 0.93; males: n = 8 per group, $F_{4,56} = 0.67$, p = 0.61).

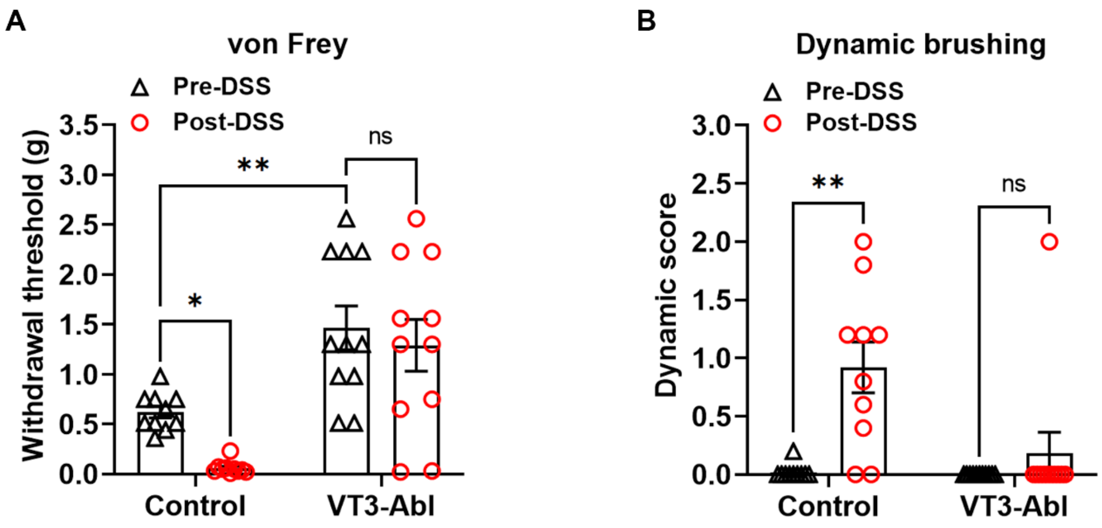


**Figure S8**. **Loss of referred cutaneous mechanical hypersensitivity in DSS-treated VGLUT3$^{LBX1}$-Abl mice**. **Related to Figure 3.**

(A) Loss of referred punctate mechanical hypersensitivity in response von Frey filament stimulation applied to the hindpaw in DSS-treated VGLUT3$^{Lbx1}$-Abl ("VT3-Abl") mice compared with control littermates (two-way ANOVA, n = 10, 11, $F_{1,38}$ = 4.14, p < 0.05; post-hoc Holm-Sidak test: *p < 0.05; ns, not significant, p = 0.41). Note that as reported previously (Cheng et al., 2017), VGLUT3$^{LBX1}$-Abl mice showed elevation of baseline withdrawal thresholds in comparison with control mice (post-hoc Holm-Sidak test: **p < 0.01). Despite this elevation of baseline thresholds, previous studies showed that withdrawal thresholds can still be reduced to the same levels seen in control mice following cutaneous inflammation or nerve lesions (Cheng et al., 2017). In other words, the lack of sensitization in DSS-treated VGLUT3$^{LBX1}$-Abl mice is not due to ceiling effects, but rather suggests requirement of VGLUT3$^{LBX1}$ neurons for the development or the expression of referred punctate mechanical hypersensitivity in DSS-treated mice.

(B) After DSS feeding, brush-evoked dynamic mechanical hypersensitivity was observed in control littermates, but not in VT3-Abl mice (two-way ANOVA, n = 10, 11, $F_{1,19}$ = 14.46, p < 0.01; post-hoc Holm-Sidak test: **p < 0.01; ns, not significant, p = 0.37).

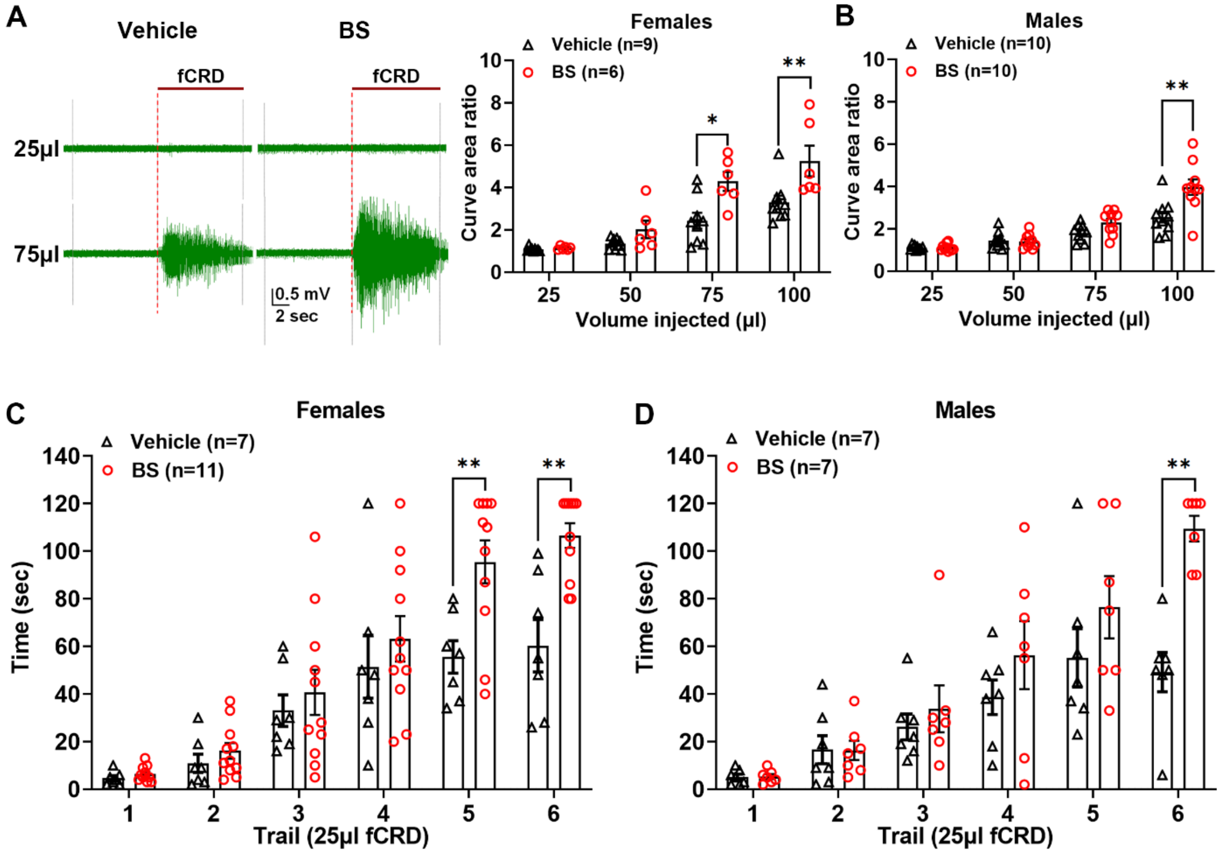


**Figure S9. Spinal disinhibition led to sensitized VMRs and aversion evoked by fCRD. Related to Figure 3.**

(A, B) Sensitized fCRD-evoked VMRs following intrathecal injection of bicuculline and strychnine (BS). Representative traces of EMG activity evoked by 25μl and 75μl fCRD (A, left). Quantitative analyses show an increase in fCRD-evoked EMG activity, for both BS-treated females and males compared with vehicle-treated mice (two-way ANOVA; females: n = 6, 9, $F_{3,39}$ = 4.27, p < 0.05; males: n = 10 per group, $F_{3,54}$ = 9.81, p < 0.01; post-hoc Holm-Sidak test: *p < 0.05, **p < 0.01). Note that the responses appeared to be stronger in females, particularly when 75μl fCRD was performed.

(C, D) 25μl fCRD evoked stronger aversive learning measured via the step-down assay in mice with intrathecal BS injection compared with vehicle injection, in both females and males (two-way ANOVA, females: n = 7, 11, $F_{5,80}$ = 3.31, p < 0.01; males: n = 7 per group, $F_{5,60}$ = 3.95, p < 0.01; post-hoc Holm-Sidak test: **p < 0.01). Note again the trend that 25μl fCRD-evoked aversive learning was stronger in females in comparison with males.

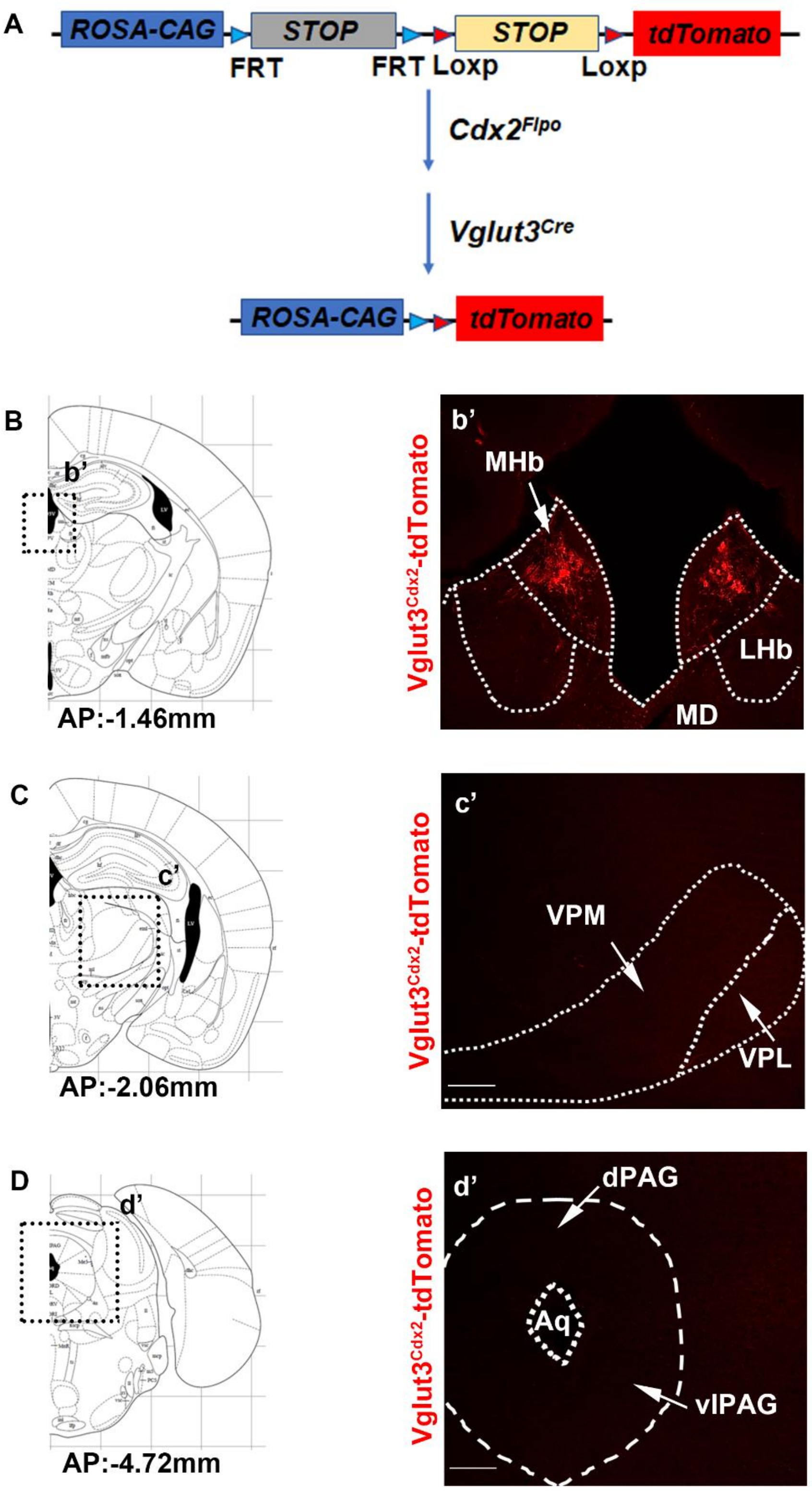


**Figure S10. Central projections of spinal VGLUT3 lineage neurons. Related to Figure 4.**

(A) Schematic illustration of the intersectional approach to produce *Vglut3$^{Cdx2}$-tdTomato* mice. Within the central nervous system, tdTomato expression was confined to VGLUT3$^{Cre}$-marked spinal neurons, upon removal of two STOP cassettes by Cre and Flpo, with Cdx2-Flpo expression confined to the spinal cord within the central nervous system.

(B-D) In *Vglut3$^{Cdx2}$-tdTomato* mice, tdTomato$^{+}$ nerve terminals derived from VGLUT3$^{Cre}$-marked spinal neurons were found in the medial habenula (B; b', mHb), but none or rarely seen in the lateral habenula (B; b', LHb), the medial thalamic nuclei (B;b', MD), the ventral posterolateral (VPL) and ventral posteromedial (VPM) thalamic nuclei (C; c'), the dorsal periaqueductal gray (dPAG) or the ventrolateral periaqueductal gray (vlPAG) (D, d'). Schematic diagrams with indicated anterior-posterior coordinates are derived from the mouse brain atlas (Franklin and Paxinos, 1997).

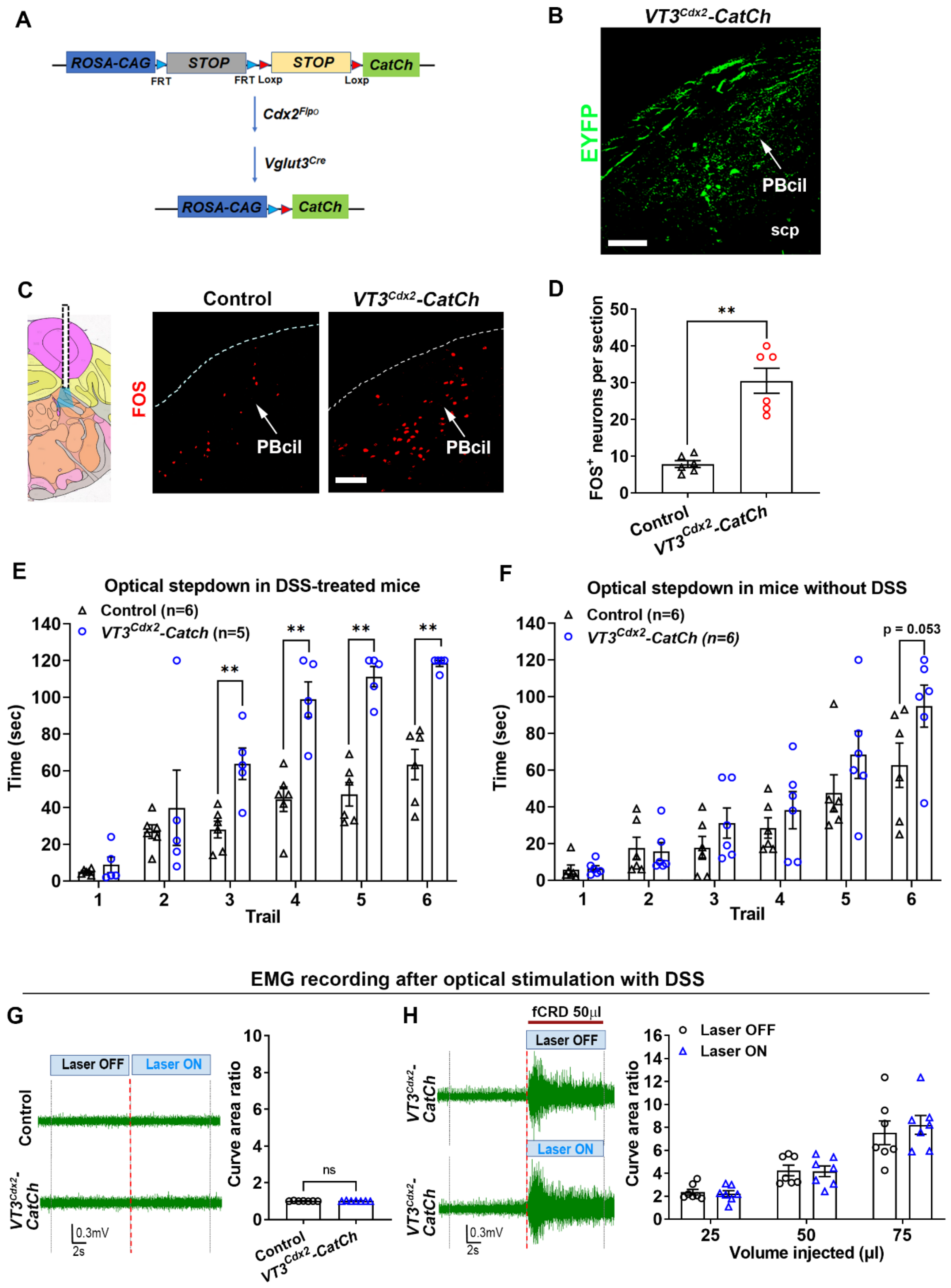


**Figure S11**. **Impact of optogenetic activation of nerve terminals originating from spinal VGLUT3 lineage neurons in caudal parabrachial nuclei**. **Related to Figure 4.**

(A) Schematic illustration of the intersectional genetic approach to create *Vglut3*$^{Cdx2}$*-CatCh* mice. CatCh is a L132C mutant channelrhodopsin with enhanced $Ca^{++}$ permeability, accelerated response kinetics and greater blue light sensitivity in comparison with the non-mutated channelrhodopsin protein (Kleinlogel et al., 2011). In these mice, expression of CatCh plus the EYFP reporter in the central nervous system was confined to VGLUT3$^{Cre}$-marked spinal neurons, upon removal of two STOP cassettes by Cre and Flpo.

(B) In the caudal PBcil of *Vglut3*$^{Cdx2}$*-CatCh* mice, nerve terminals bearing the EYFP reporter signal were observed following anti-GFP immunostaining.

(C, D) 473nm blue laser stimulation of CatCh$^{+}$ terminals in caudal PBcil (according to the mouse brain atlas coordinates: AP: -5.3 mm; ML: -1.3 mm; DV: -2.3mm) was able to increase FOS$^{+}$ neurons per PBcil section from *Vglut3*$^{Cdx2}$*-CatCh* mice compared with control littermates (Student's *t*-test, n = 6 per group, $t_{10}$ = 6.38, **p < 0.01).

(E) 15 Hz optical stimulation of terminals in the caudal PBcil region drove aversive learning in DSS-treated *Vglut3*$^{Cdx2}$*-CatCh* mice compared with control littermates (two-way ANOVA, n = 5, 6, $F_{5,45}$ = 5.41, p < 0.01; post-hoc Holm-Sidak test: **p < 0.01). This aversive learning was comparable to the learning produced by 30 Hz stimulation shown in Figure 4F. Data are shown as mean ± SEM.

(F) 30 Hz optical stimulation in the caudal PBcil region induced a trend of aversive learning in naïve *Vglut3*$^{Cdx2}$*-CatCh* mice compared with control littermates (two-way ANOVA, n = 6 per group, $F_{5,50}$ = 1.29, p = 0.28; post-hoc Holm-Sidak test: p = 0.053).

(G) Optical stimulation of CatCh$^{+}$ terminals in the caudal PBcil did not induce any EMG responses. Representative EMG traces with 473nm blue laser stimulation (G, left), and no difference between control mice and *Vglut3*$^{Cdx2}$*-CatCh* mice (G, right, Student's *t*-test, n = 7 per group, $t_{12}$ = 0.03; ns, not significant, p = 0.97).

(H) Optical stimulation of CatCh$^{+}$ terminals in the caudal PBcil failed to enhance fCRD-evoked VMRs in DSS-treated *Vglut3*$^{Cdx2}$*-Catch* mice compared with control littermates. Representative EMG traces evoked by 50 μl fCRD with or without 473nm blue laser stimulation, and no difference was detected (two-way ANOVA, n = 7 per group, $F_{2,18}$ = 0.95, p = 0.41).
